# A multi-year longitudinal harmonization of quality controls in mass spectrometry proteomics core facilities

**DOI:** 10.1101/2024.05.19.594517

**Authors:** Cristina Chiva, Roger Olivella, An Staes, Teresa Mendes Maia, Christian Panse, Karel Stejskal, Thibaut Douché, Bérangère Lombard, Andrea Schuhmann, Damarys Loew, Karl Mechtler, Mariette Matondo, Mandy Rettel, Dominic Helm, Francis Impens, Simon Devos, Anna Shevchenko, Paolo Nanni, Eduard Sabidó

## Abstract

Quality control procedures play a pivotal role in ensuring the reliability and consistency of data generated in mass spectrometry-based proteomics laboratories. However, the lack of standardized quality control practices across laboratories poses challenges for data comparability and reproducibility. In response, we conducted a harmonization study within proteomics laboratories of the Core for Life alliance with the aim to establish a common quality control framework, which facilitates comprehensive quality assessment and identification of potential sources of performance drift. Through collaborative efforts, we developed a consensus quality control standard for longitudinal assessment and we adopted a common processing software. We generated a 4-year longitudinal dataset from multiple instruments and laboratories, which enabled us to assess intra- and inter-laboratory variability, to identify causes of performance drift, and to establish community reference values for several quality control parameters. Our study enhances data comparability and reliability and fosters a culture of collaboration and continuous improvement within the proteomics community to ensure the integrity of proteomics data.

## Introduction

Common quality control procedures in proteomics core facilities ensure technical quality, reproducibility, comparability, and integrity of proteomics data, thus resulting in multiple benefits for the proteomics infrastructures, the scientific community and the users and customers of these infrastructures. As representatives of eight European proteomics research infrastructures and core facilities integrated within the Core for Life alliance (Meder et al, 2016; Lippens et al, 2019) we have recently advocated for community policy for quality control procedures to ensure the high standards in proteomics services (Chiva et al, 2021). The presence of such common quality procedures is especially relevant as core facilities are increasingly becoming key players in research projects. Despite their importance, the current landscape reveals a general lack of common quality control procedures across proteomics laboratories, which employ a diverse range of protocols, standards, reference compounds, and software tools. There is therefore a need for a common framework and set of procedures that can facilitate universal quality control and assessment among independent proteomics laboratories, while remaining flexible to their varied instrumentation and setups.

Based on our previous work, here we set out to create a common quality control framework specifically designed for mass spectrometry-based proteomics laboratories with the goal to identify and reduce potential sources of performance drift, ensuring the reliability and consistency of data generated in various experimental setups. In line with this objective we initiated a harmonization study among the proteomics core facilitates within the Core for Life alliance (https://coreforlife.eu) to improve the comparability of the quality control results and thus identify community reference values, intra-laboratory performance drifts, potential sources of heterogeneity, and opportunities for improvement. In contrast to standardization, harmonization studies do not impose homogeneous methods, but a general agreement on a series of practices for prospective data collection (Fortier et al, 2017). Indeed, imposing identical methodologies has demonstrated to be challenging in research environments due to the different available equipment, diverse setups, and different scientific interests and priorities that exist in proteomics core facilities and research infrastructures. Harmonization frameworks are therefore more flexible, and they embrace a broader range of setups and instrumentation, overcoming the difficulties of many standardization initiatives.

As a result of our harmonization study, we successfully established a common quality control framework within the mass spectrometry-based proteomics laboratories of the Core for Life alliance. This framework facilitated comprehensive quality control discussions, comparisons, and the identification and mitigation of potential sources of performance drift. Moreover, as a tangible outcome of our efforts, we report here i) a new consensus quality control standard for longitudinal assessment of liquid chromatography and mass spectrometry system (LC-MS) performance in proteomics setups, ii) a common quality control processing software, iii) measures of intra- and inter-laboratory variability conducted in between real sample analyses, iv) a set of community reference values for several quality control parameters, and v) 4-year longitudinal series of quality control data from multiple instruments and laboratories.

## Experimental Methods

In our harmonization study, we established standardized parameters for each instrument type to measure a common quality control sample, while allowing flexibility in the HPLC settings to accommodate the diverse analytical setups across sites. While we acknowledge that this approach may introduce variations among sites and instruments, our intention was to operate within real-world conditions and strive for harmonization rather than strict standardization.

### Orbitrap Q Exactive HF

#### Common parameters

25 ng of QC4L digest were analyzed in Orbitrap Q Exactive HF. The Orbitrap Q Exactive HF mass spectrometer was operated in data-dependent performing a full scan (m/z range 350-1500, resolution 120,000, target value 3E6), followed by MS/MS scans of the 12 most abundant ions. MS/MS spectra were acquired using an isolation width of 1.2 m/z, target value of 1E5 and intensity threshold of 9E4, maximum injection time 50 ms, HCD collision energy of 28. Precursor ions selected for fragmentation (include charge state 2-6) were excluded for 30 s. The monoisotopic precursor selection filter and exclude isotopes feature were enabled. Up-to-date Xcalibur tune versions were used during the harmonization study.

#### Specific parameters

##### Flanders Institute for Biotechnology (Site 5)

The UltiMate 3000 RSLC nano system (Thermo Fisher Scientific) or the Evosep One LC system (Evosep) was used coupled to one of the two available Orbitrap Q Exactive HF mass spectrometers. Peptides were loaded onto a trap column (in-house C18 column, Reprosil-HD Dr. Maisch, 20 mm × 100 μm ID, 5 μm) with loading buffer A (2:98 acetonitrile:water, 0.1% trifluoracetic acid) at a flow rate of 10 µL/min over 4 min from April 2019 to Dec 2021, whereas from Jan 2022 onwards loading was performed onto a new trap column (PepMap Acclaim C18, 5 mm × 300 μm ID, 5 μm, 100 Å, Thermo Fisher Scientific) at a flow rate of 20 µL/min over 2 min. Separation was then performed on the analytical column (200-cm µPAC Gen1 C18 column endcapped functionality April 2019-April 2020, operated at 50°C with a linear gradient from 0 to 33% buffer B (20:80 water:acetonitril, 0.1% formic acid) in 42 min, and reaching 55% buffer B at 58 min. Buffer A: 0.1% formic acid in water. The first 15 min the flow rate was set to 750 nl/min and then lowered to 230 nl/min. From May 2020 until Oct 2020, in the course of Feb 2021 until first half of July 2021, and again from Nov 2021 on until the first half of March 2022 separation was performed on a 25-cm analytical column (nanoEase M/Z HSS T3 Column, 250 mm x 75 µm ID, 1.8 µm, 100 Å, Waters). From March 2022 until May 2022 separation was performed on an in-house 25-cm analytical column (in-house C18 column, Reprosil-HD Dr. Maisch, 250 mm x 75 μm ID, 1.9 μm), and starting from June 2022 until the end of 2022 separation was performed with a commercial analytical column (Aurora^TM^ Ultimate 25cm column, 250 mm x 75 μm ID, 1.9 μm, 120 Å, IonOptiks). All columns were kept at a constant temperature of 45°C either in the Ultimate 3000 column compartment or in a butterfly oven (Phoenix Science & Technology Inc). The Evosep One LC system was connected to the mass spectrometer from Oct 2020 until Jan 2021, from July 2021 until Oct 2021, and between June and Aug 2022. During the first and second period 100 ng of QC4L digest was loaded on the Evotip, while during the last period 25 ng of QC4L digest was loaded. In the first two periods the peptides were analyzed with the 15 SPD method using the endurance Evosep column (150 mm x 150 µm, 1.9 µm) whereas the performance Evosep column (150 mm x 75 µm, 1.9 µm) with the 20 SPD whisper method was used for the last period. MS settings were kept constant across the 4 years of the study aligned with the other institutes.

The UltiMate 3000 RSLC nano system (Thermo Fisher Scientific) was used coupled to the second available Orbitrap Q Exactive HF mass spectrometer. A dual column setup was used, consisting of two nano-pumps and one loading pump. Peptides were loaded onto a trap column (in-house C18 column, Reprosil-HD Dr. Maisch, 20 mm × 100 μm ID, 5 μm) with loading buffer A (2:98 acetonitrile:water, 0.1% trifluoracetic acid) at a flow rate of 10 µL/min over 4 min from Jan 2019 to June 2022, whereas from July 2022 onwards loading was performed onto a new trap column (PepMap Acclaim C18, 5 mm × 300 μm ID, 5 μm, 100 Å, Thermo Fisher Scientific) at a flow rate of 20 µL/min over 2 min. On both flowpaths, separation was then performed on the analytical column (200-cm µPAC Gen1 C18 column endcapped functionality April 2019-June 2022, and alternating 110-cm µPAC Gen2 C18 column July 2022 onwards, Thermo Fisher Scientific) operated at 50°C. On 200-cm µPAC Gen1 C18 columns peptides were eluted with a linear gradient from 0-33 % buffer B (80% acetonitrile, 0.1% formic acid) in 42 min, and 55% at 58 min. On 110-cm µPAC Gen2 C18 columns peptides were eluted by a linear gradient from 0 to 26.4% buffer B (100% acetonitrile, 0.1% formic acid) in 60 min, and 44% at 68 min. During the first 15 min the flow rate was set to 750 nl/min for the 200-cm µPAC Gen1 C18 column, and to 600 nl/min flow rate for 5 min for the 110-cm µPAC Gen2 C18 column after which the flow rate for these two types of columns was kept constant at 300 nl/min. A Pneu-Nimbus source (Phoenix Science & Technology Inc) was used for dual column setup, switching between the two column setups between every run.

##### Functional Genomics Center Zurich (Site 3)

The ACQUITY UPLC M-Class system (Waters) was used coupled to the mass spectrometer, with a Digital PicoView source (New Objective). Peptides were loaded on a trap column (ACQUITY UPLC M-Class Symmetry C18, 20 mm x 180 µm ID, 5 μm particles, 100 Å pore size, Waters) with buffer A (100% water, 0.1% formic acid). Separation was then performed on the analytical column (nanoEase MZ C18 HSS T3 Column, 250 mm x 75 µm, 1.8 μm, 100 Å, Waters) operated at 50 °C with a linear gradient from 8% to 30% buffer B (100% acetonitrile, 0.1% formic acid) at a flow rate of 300 nl/min over 60 min.

##### Institut Pasteur (Site 7)

EASY-nLC 1200 system (Thermo Scientific) was used coupled to the mass spectrometer, with an integrated column oven (PRSO-V1 - Sonation GmbH, Biberach, germany). Peptides were directly loaded into analytical C18 columns (in-house, ReproSil-Pur Basic C18 Dr. Maisch GmbH, 200-500 mm x 75 µm ID, 1.9 µm, 100 Å) with buffer A (100% water, 0.1% formic acid) at a constant pressure of 900 bar and 40 °C. Separation was performed with a multi-step gradient from 2 to 7% buffer B (80% acetonitrile, 0.1% FA) in 1 min, to 23% in 45 min, and to 45% in 9 min at a flow rate of 250 nL/min.

##### Max Planck Institute of Molecular Cell Biology and Genetics (Site 6)

The Ultimate 3000 RSLC nano LC system (Thermo Scientific) was coupled to the mass spectrometer. The LC system was equipped with Acclaim™ PepMap™ 100 C18 trapping column (20 mm × 100 μm ID, 5 μm, 100 Å, Thermo Scientific) and a variety of analytical columns: before September 2020/October 2020-February 2021 – with Acclaim™ PepMap™ C18 (150 mm × 75 μm ID, 3 μm, 100), September 2020 – with Gen1 µPAC 200 cm, from February 2021 – with Gen1 µPAC 50 cm (all Thermo Scientific). All columns were kept at constant temperature of 50°C. Peptides were loaded at 6 μl/min flow rate in solvent A (0.1% aquatic formic acid) and eluted at 500 nl/min (Gen1 µPAC) or 300 nl/min (Acclaim™ PepMap™) by 60 min multi-step linear gradient as following: 0% to 30% solvent B (0.1% formic acid in acetonitrile) 0-45 min, 30% to 50% B at 46-58 min and 100% B 58-60 min. The column was then washed with B for 5 minutes and re-equilibration with solvent A for 15 min. Peptides were introduced into the Q Exactive HF using nano ESI Emitters 360 µm OD x 20 µm ID, 20 µm tip (Fossiliontech). The Nanospray Flex™ Source (Thermo Scientific) spray voltage was set on 2.5 kV. The capillary temperature was set at 280°C.

### Orbitrap Q Exactive HF-X

#### Common parameters

25 ng of QC4L digest were analyzed in Orbitrap Q Exactive HF-X. The Orbitrap Q Exactive HF-X mass spectrometer was operated in data-dependent performing a full scan (m/z range 350-1500, resolution 120,000, target value 3E6), followed by MS/MS scans of the 12 most abundant ions. MS/MS spectra were acquired using an isolation width of 1.2 m/z, target value of 1E5 and intensity threshold of 9E4, maximum injection time 50 ms, HCD collision energy of 28. Precursor ions selected for fragmentation (include charge state 2-6) were excluded for 30 s. The monoisotopic precursor selection filter and exclude isotopes feature were enabled. Up-to-date Xcalibur tune versions were used during the harmonization study.

#### Specific parameters

##### Functional Genomics Center Zurich (Site 3)

The ACQUITY UPLC M-Class system (Waters) was used coupled to the mass spectrometer, with a Digital PicoView source (New Objective). Peptides were loaded on a trap column (ACQUITY UPLC M-Class Symmetry C18, 20 mm x 180 µm ID, 5 μm particles, 100 Å pore size, Waters) with buffer A (100% water, 0.1% formic acid). Separation was then performed on the analytical column (nanoEase MZ C18 HSS T3 Column, 250 mm x 75 µm, 1.8 μm, 100 Å, Waters) operated at 50 °C with a linear gradient from 8% to 30% buffer B (100% acetonitrile, 0.1% formic acid) at a flow rate of 300 nl/min over 60 min.

##### Institut Curie (Site 8)

The UltiMate 3000 RSLC nano system (Thermo Fisher Scientific) was used coupled to the mass spectrometer, with a Nanosrpay Flex ion source (Thermo Fisher Scientific). Peptides were first trapped onto a C18 column (PepMap Acclaim C18, 20 mm x 75 μm ID; 5 μm, 100 Å, Thermo Fisher Scientific) with buffer A (100% water, 0.1% formic acid) at a flow rate of 2.5 µL/min over 4 min. Separation was then performed on the analytical column (PepMap Acclaim C18, 500 mm × 75 μm ID, 2 μm, 100 Å, Thermo Fisher Scientific) operated at 50 °C with a linear gradient from 2% to 30% buffer B (100% acetonitrile, 0.1% formic acid) at a flow rate of 300 nL/min over 60 min.

##### The Research Institute of Molecular Pathology (Site 4)

The UltiMate 3000 RSLC nano system (Thermo Fisher Scientific) was used coupled to the mass spectrometer, with a Nanospray Flex ion source (Thermo Fisher Scientific). Peptides were loaded onto a trap column (PepMap Acclaim C18, 5 mm × 300 μm ID, 5 μm, 100 Å, Thermo Fisher Scientific) with loading buffer (100% water, 0.1% trifluoracetic acid) at a flow rate of 25 µL/min over 10 min. Separation was then performed on the analytical column (PepMap Acclaim C18, 500 mm × 75 μm ID, 2 μm, 100 Å, Thermo Fisher Scientific) operated at 30°C with a linear gradient from 2% to 35% buffer B (80% acetonitrile, 0.1% formic acid) at a flow rate of 230 nl/min over 60 min. Buffer A: 100% water, 0.1% formic acid.

### Orbitrap Fusion Lumos

#### Common parameters

25 ng of QC4L digest were analyzed in Orbitrap Fusion Lumos. The Orbitrap Fusion Lumos mass spectrometer was operated in data-dependent, and following each a full scan (m/z range 350-1500, resolution 120,000, target value 1E5, max injection time 50 ms), followed the 2 most intense precursors were selected for further fragmentation (charge state 2-7, intensity threshold 5E3, dynamic exclusion 60 s). Fragmentation spectra were acquired using four different acquisition settings: i) ddMS2 IT HCD with an isolation window of 1.6 m/z, collision energy of 28, maximum injection time 200 ms, AGC target standard and rapid scan rate; ii) ddMS2 IT CID with an isolation window of 1.6 m/z, collision energy of 35, max injection time 200 ms, AGC target standard and rapid scan rate; iii) ddMS2 IT ETciD with an isolation window of 1.6 m/z, collision energy of 35, max injection time 200 ms, AGC target 110% and rapid scan rate; and iv) ddMS2 IT EThcD with an isolation window of 1.6 m/z, collision energy of 15, max injection time 200 ms, and AGC target 110% and rapid scan rate. Up-to-date Xcalibur tune versions were used during the harmonization study.

#### Specific parameters

##### Centre for Genomic Regulation (Site 1)

The EASY-nLC 1200 system (Thermo Fisher Scientific) was used coupled to the mass spectrometer, with a EASY-Spray ion source (Thermo Fisher Scientific). Peptides were loaded directly onto the analytical column (EASY-Spray C18 column, 500 mm × 75 μm ID, 2 μm, 100 Å, Thermo Fisher Scientific) operated at 45°C with buffer A (100% water, 0.1% formic acid) at a flow rate of 300 nL/min. Separation was then performed with a linear gradient from 5% to 25% buffer B (80% acetonitrile, 0.1% formic acid) over 52 min and up 40 % buffer B over 8 min at a flow rate of 300 nl/min.

##### European Molecular Biology Laboratory (Site 2)

The UltiMate 3000 RSLC nano system (Thermo Fisher Scientific) was used coupled to the mass spectrometer, with a Nanospray Flex ion source (Thermo Fisher Scientific). Peptides were loaded onto a trap column (PepMap Acclaim C18, 5 mm × 300 μm ID, 5 μm, 100 Å, Thermo Fisher Scientific) with loading buffer (100% water, 0.05% trifluoracetic acid) at a flow rate of 30 µL/min over 4 min. Separation was then performed on the analytical column (nanoEase M/Z HSS T3 C18 column, 250 mm × 75 μm ID, 1.8 μm, 100 Å, Waters) operated at 40°C with a linear gradient from 2% to 40% buffer B (3% DMSO, 0.1% formic acid in acetonitrile) at a flow rate of 300 nl/min over 70 min. Buffer A: 3% DMSO, 0.1% formic acid in water.

##### Flanders Institute for Biotechnology (Site 5)

The UltiMate 3000 RSLC nano system (Thermo Fisher Scientific) was used coupled to the mass spectrometer, with a Nanospray Flex ion source (Thermo Fisher Scientific). A dual column setup was used, consisting of two nano-pumps and one loading pump. Peptides were loaded onto a trap column (in-house C18 column, Reprosil-HD Dr. Maisch, 20 mm × 100 μm ID, 5 μm) with loading buffer A (2:98 acetonitrile:water, 0.1% trifluoracetic acid) at a flow rate of 10 µL/min over 4 min from April 2019 to Dec 2021, whereas from Jan 2022 onwards loading was performed onto a new trap column (PepMap Acclaim C18, 5 mm × 300 μm ID, 5 μm, 100 Å, Thermo Fisher Scientific) at a flow rate of 20 µL/min over 2 min. Separation was then performed on the analytical column (200-cm µPAC Gen1 C18 column endcapped functionality April 2019-Dec 2021, and alternating 110-cm and 50-cm µPAC Gen2 C18 column Jan 2022 onwards, Thermo Fisher Scientific) operated at 50°C with a linear gradient from 2% to 35% buffer B (80% acetonitrile, 0.1% formic acid) at a flow rate of 230 nl/min over 60 min. Peptides were eluted by a linear gradient on 200-cm µPAC Gen1 C18 columns from 0 to 33% buffer B (80% acetonitrile, 0.1% formic acid) in 42 min, and 55% at 58 min. On 50-cm µPAC Gen2 C18 columns peptides were eluted by a linear gradient from 0 to 22.5% buffer B (100% acetonitrile, 0.1% formic acid) in 55 min, and 55% at 75 min. On 110-cm µPAC Gen2 C18 columns peptides were eluted by a linear gradient from 0 to 26.4% buffer B (100% acetonitrile, 0.1% formic acid) in 60 min, and 44% at 68 min. During the first 15 min the flow rate was set to 750 nl/min for the 200-cm µPAC Gen1 C18 column, and to 600 nl/min flow rate for 5 min for the 110-cm µPAC Gen2 C18 column after which the flow rate for these two types of columns was kept constant at 300 nl/min. The flow rate of the 50-cm µPAC Gen2 columns was kept at 250 nl/min at all times. A Pneu-Nimbus source (Phoenix Science & Technology Inc) was used for dual column setup, switching between the two column setups between every run.

##### Functional Genomics Center Zurich (Site 3)

The ACQUITY UPLC M-Class system (Waters) was used coupled to the mass spectrometer, with a Digital PicoView source (New Objective). Peptides were loaded on a trap column (ACQUITY UPLC M-Class Symmetry C18, 20 mm x 180 µm ID, 5 μm particles, 100 Å pore size, Waters) with buffer A (100% water, 0.1% formic acid). Separation was then performed on the analytical column (ACQUITY UPLC M-Class C18 HSS T3 Column, 250 mm x 75 µm, 1.8 μm, 100 Å, Waters) operated at 50 °C with a linear gradient from 8% to 22% buffer B (100% acetonitrile, 0.1% formic acid) over 52 min and up 32 % buffer B over 8 min at a flow rate of 300 nl/min.

##### The Research Institute of Molecular Pathology (Site 4)

The UltiMate 3000 RSLC nano system (Thermo Fisher Scientific) was used coupled to the mass spectrometer, with a Nanospray Flex ion source (Thermo Fisher Scientific). Peptides were loaded onto a trap column (PepMap Acclaim C18, 5 mm × 300 μm ID, 5 μm, 100 Å, Thermo Fisher Scientific) with loading buffer (100% water, 0.1% trifluoracetic acid) at a flow rate of 10 µL/min over 10 min. Separation was then performed on the analytical column (PepMap Acclaim C18, 500 mm × 75 μm ID, 2 μm, 100 Å, Thermo Fisher Scientific) operated at 30°C with a linear gradient from 2% to 35% buffer B (80% acetonitrile, 0.1% formic acid) at a flow rate of 230 nl/min over 30 min. Buffer A: 100% water, 0.1% formic acid.

### Orbitrap Eclipse

#### Common parameters

25 ng of QC4L digest were analyzed in Orbitrap Eclipse. The Orbitrap Fusion Lumos mass spectrometer was operated in data-dependent, and following each a full scan (m/z range 350-1500, resolution 120,000, target value 1E5, max injection time 50 ms), followed the 2 most intense precursors were selected for further fragmentation (charge state 2-7, intensity threshold 5E3, dynamic exclusion 60 s). Fragmentation spectra were acquired using four different acquisition settings: i) ddMS2 IT HCD with an isolation window of 1.6 m/z, collision energy of 28, maximum injection time 200 ms, AGC target standard and rapid scan rate; ii) ddMS2 IT CID with an isolation window of 1.6 m/z, collision energy of 35, max injection time 200 ms, AGC target standard and rapid scan rate; iii) ddMS2 IT ETciD with an isolation window of 1.6 m/z, collision energy of 35, max injection time 200 ms, AGC target 110% and rapid scan rate; and iv) ddMS2 IT EThcD with an isolation window of 1.6 m/z, collision energy of 15, max injection time 200 ms, and AGC target 110% and rapid scan rate. Up-to-date Xcalibur tune versions were used during the harmonization study.

#### Specific parameters

##### Centre for Genomic Regulation (Site 1)

The EASY-nLC 1200 system (Thermo Fisher Scientific) was used coupled to the mass spectrometer, with a EASY-Spray ion source (Thermo Fisher Scientific). Peptides were loaded directly onto the analytical column (EASY-Spray C18 column, 500 mm × 75 μm ID, 2 μm, 100 Å, Thermo Fisher Scientific) operated at 45°C with buffer A (100% water, 0.1% formic acid) at a flow rate of 300 nL/min. Separation was then performed with a linear gradient from 5% to 25% buffer B (80% acetonitrile, 0.1% formic acid) over 52 min and up 40 % buffer B over 8 min at a flow rate of 300 nl/min.

### Orbitrap Exploris 480

#### Common parameters

12.5 ng of QC4L digest were analyzed in Orbitrap Exploris 480. The Orbitrap Exploris 480 mass spectrometer was operated in data-dependent performing a full scan (m/z range 380-1200, resolution 60,000, AGC target 300%), followed by MS/MS scans of the most abundant ions. MS/MS spectra were acquired using an isolation width of 0.7 m/z, AGC 100% and intensity threshold of 1E4, maximum injection time set to auto, HCD collision energy of 30. Precursor ions selected for fragmentation (include charge state 2-6) were excluded for 20 s. The monoisotopic precursor selection filter and exclude isotopes feature were enabled. Up-to-date Xcalibur tune versions were used during the harmonization study.

#### Specific parameters

##### Functional Genomics Center Zurich (Site 3)

The ACQUITY UPLC M-Class system (Waters) was used coupled to the mass spectrometer, with a Digital PicoView source (New Objective) and a Nanospray Flex Ion (Thermo Fisher Scientific). Peptides were loaded on a trap column (ACQUITY UPLC M-Class Symmetry C18, 20 mm x 180 µm ID, 5 μm particles, 100 Å pore size, Waters) with buffer A (100% water, 0.1% formic acid). Separation was then performed on the analytical column (ACQUITY UPLC M-Class C18 HSS T3 Column, 250 mm x 75 µm, 1.8 μm, 100 Å, Waters) operated at 50 °C with a linear gradient from 5% to 35% buffer B (100% acetonitrile, 0.1% formic acid) at a flow rate of 300 nl/min over 30 min.

##### Institut Curie (Site 8)

The UltiMate 3000 RSLC nano system (Thermo Fisher Scientific) was used coupled to the mass spectrometer, with a Nanospray Flex ion source (Thermo Fisher Scientific). Peptides were first trapped onto a C18 column (PepMap Acclaim C18, 20 mm x 75 μm ID; 5 μm, 100 Å, Thermo Fisher Scientific) with buffer A (2:98 acetonitrile:water, 0.1% formic acid) at a flow rate of 3 µL/min over 4 min. Separation was then performed on the analytical column (PepMap Acclaim C18, 500 mm × 75 μm ID, 3 μm, 100 Å, Thermo Fisher Scientific) operated at 40 °C with a linear gradient from 3% to 30% buffer B (100% acetonitrile, 0.1% formic acid) at a flow rate of 300 nL/min over 30 min.

##### The Research Institute of Molecular Pathology (Site 4)

The UltiMate 3000 RSLC nano system (Thermo Fisher Scientific) was used coupled to the mass spectrometer, with a Nanospray Flex ion source (Thermo Fisher Scientific). Peptides were loaded onto a trap column (PepMap Acclaim C18, 5 mm × 300 μm ID, 5 μm, 100 Å, Thermo Fisher Scientific) with loading buffer (100% water, 0.1% trifluoracetic acid) at a flow rate of 10 µL/min over 10 min. Separation was then performed on the analytical column (PepMap Acclaim C18, 500 mm × 75 μm ID, 2 μm, 100 Å, Thermo Fisher Scientific) operated at 30°C with a linear gradient from 2% to 35% buffer B (80% acetonitrile, 0.1% formic acid) at a flow rate of 230 nl/min over 30 min. Buffer A: 100% water, 0.1% formic acid.

### Data analysis

The analysis of all the raw files and the extraction of the quality control parameters has been done through the standard pipeline available in the Qcloud platform (Chiva et al. 2018, Olivella et al. 2021).

## Results and discussion

Often quality control and quality assessment procedures in proteomics core facilities and research infrastructures present a motley collection of protocols, standards, reference compounds and software tools. Therefore, as a first step to the creation of a common framework for quality control assessment, we set to define common standards and software tools, and a comprehensive array of parameters to monitor as a first step in our harmonization study to improve the comparability among quality control results. At the same time, however, being aware of the heterogeneity of setups, we allowed some flexibility to embrace a broader range of configurations that reflect real day-to-day instrument usage, and thus avoid artificial setups exclusively dedicated to the harmonization study.

### A new common quality control standard

Firstly, we established a common quality control standard as a way to homogenize and set a common ground towards the comparability of quality control parameters. We looked for a sample that would enable us to combine both peptide identification and quantitative parameters in one run, and that at the same time could be used as quality control and system suitability test sample, i.e. a combination of the QC1 and QC2 concepts according to the nomenclature established by Pichler et al. back in 2012 (Pichler et al, 2012). In 2015 a novel standard was described consisting of six peptides, each of them, in five different isotopologue variants and each in a different amount (Beri et al, 2015). The six peptides were designed to cover a wide range of hydrophobicities and by combining these isotopologues at different ratios, they spanned four orders of magnitude within each distinct peptide sequence. Based on this standard, we designed a new mix consisting of 25 ng K562 cells proteome extract with the isotopologue mix spiked-in covering five different concentrations ranging from 0.01 to 100 fmol, which is now commercially available through a commercial partner (QC4Life Standard Mix, Promega, CS30240). Small proteome quantities (12.5 - 25 ng) provide a complex mix to assess instrument sensitivity, and at the same time avoid carryover that might influence subsequent samples. Moreover, the presence of the peptide isotopologue mix ensures the presence of known ratios among different isotopically labeled variants of the same peptide covering four orders of magnitude, thus enabling the assessment of quantitative measurements and linear range of quantification at chromatographic times during the analysis.

### Common quality control processing software

We also established a common software platform to evaluate the results and thus minimize data analysis heterogeneity. QCloud has facilitated automated quality control of LC-MS instrumentation for many individual laboratories for the last few years (Chiva et al, 2018). Importantly, its most recent version nowadays supports several dozens of instruments from multiple laboratories worldwide (Olivella et al, 2021). We therefore adopted this software for automated, reproducible, and centralized analysis of quality control raw files generated by the different instruments in each of the participating sites. For this particular harmonization study, we adapted the QCloud2 pipeline to support QC4Life Standard Mix and we have incorporated the new signal extraction algorithm from rawDiag (Trachsel et al, 2018). Moreover, additional control charts showing the intensity of the different isotopologues over time have been incorporated and a single-file view showing the intensity of the six isotopologue peptides versus the five different concentrations are now shown. This view assists in evaluating the dynamic range for each standard peptide within the QC4Life Standard Mix (Supplementary Figure S1 and S2). A total of 109 quality control parameters were systematically extracted from each result file across various instrument types throughout this study. These parameters comprised metrics such as mass accuracy, retention time, and peak area values for each spiked-in isotopologue peptide, as well as encompassing total ion current (TIC), median IT MS1 and MS2 spectra, and counts of MS2 spectra, PSMs, uniquely identified peptides, and proteins for each available fragmentation type (e.g., CID, HCD, ETciD, EThcD) (Supplementary Table ST1). These quality control parameters were collectively and meticulously selected to evaluate the diverse aspects of LC-MS/MS performance. Notably, metrics like the number of detected peptides or MS/MS spectra facilitate rapid identification of major system malfunctions necessitating immediate intervention, while other parameters, such as drifts in injection times, or dynamic range sensitivity observed in the spiked-in isotopologue peptides, may guide fine-tuning of system adjustments or prompt further investigative tests. All quality control values were automatically extracted by the QCloud pipeline.

### Harmonization study

#### Study design

Eight different proteomics reference core facilities (CRG, EMBL, FGCZ, Institut Curie, IMP, Institut Pasteur, MPI-CBG, and VIB) have incorporated 30 different LC-MS systems corresponding to 5 different types of mass spectrometer instruments (Orbitrap Eclipse, Orbitrap Fusion Lumos, Orbitrap Q Exactive HF, Orbitrap Q Exactive HF-X, and Orbitrap Exploris 480) and 4 different liquid chromatography systems (UltiMate 3000 RSLC, EASY-nLC 1200, ACQUITY UPLC M-Class, Evosep One) into an harmonization study that spanned for a total of four years (2019-2022). The QC4Life standard sample has regularly been analyzed with each LC-MS setup using a shotgun data-dependent acquisition (DDA) method with HCD fragmentation (and ETD, EThcD and CID variants when available) and a flexible chromatographic H_2_O:ACN gradient lengths from 30-60 min. A total of 9054 raw files have been generated and 109 parameters have been extracted for each file, making a total of 826,696 quality control data points (Figure 1, Supplementary Table ST1 and ST2). During the study each laboratory has had access to its own data through QCloud, as well as the data of the rest of the sites participating in this study, which was shared among the participants through a weekly report (Supplementary Information).

**Figure 1:**
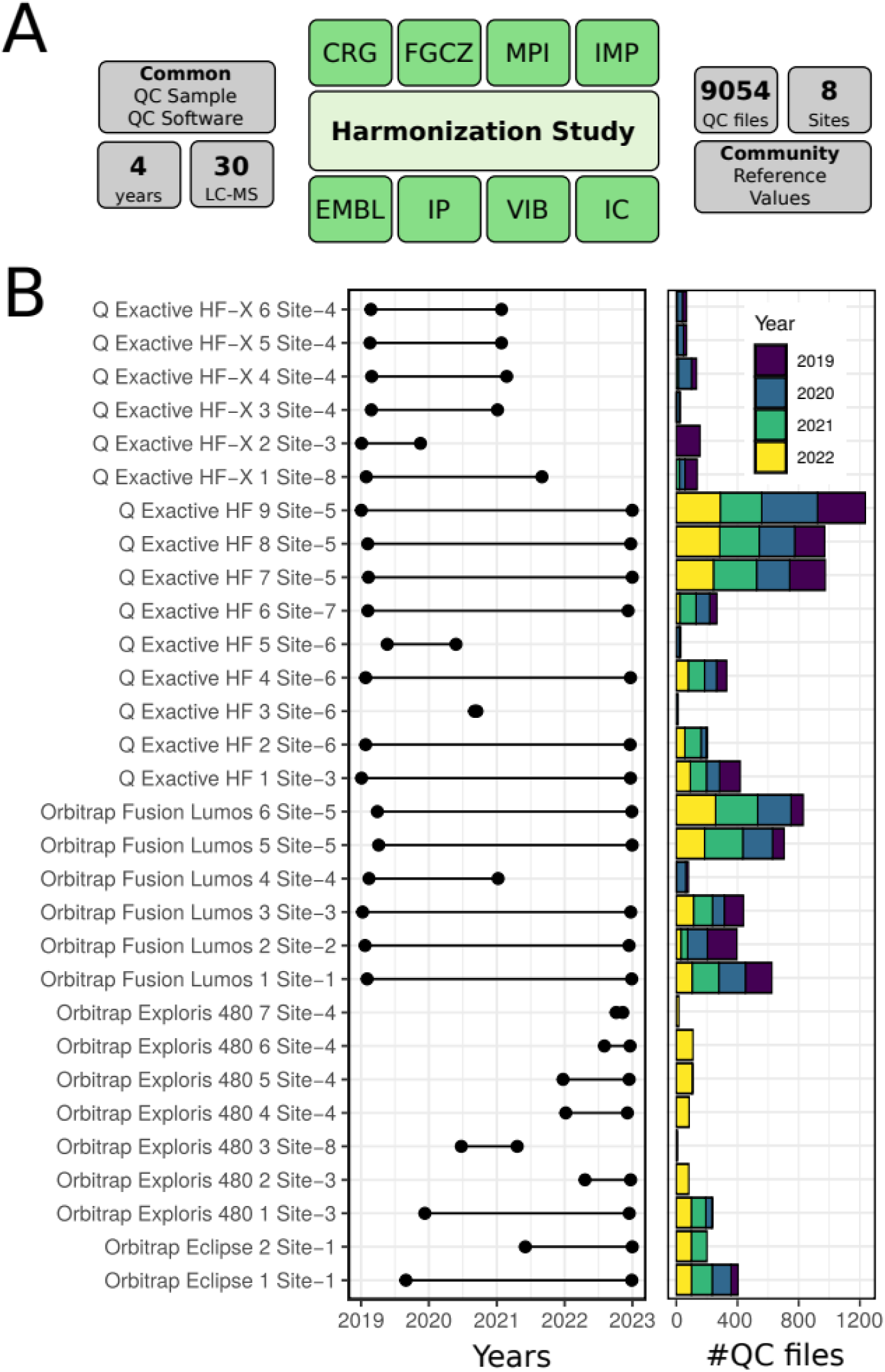
Overview of the harmonization study conducted by the Core for Life Proteomics Working group. Eight different sites have incorporated 30 different LC-MS systems corresponding to 5 different mass spectrometer instrument types and 4 liquid chromatography types into a 4-year harmonization study using a common quality control sample and a common software tool. A total of 9054 quality control raw files have been generated and 109 quality control parameters have been extracted for each file, making a total of 826,696 quality control data points.

#### Measure of intra- and inter-laboratory variability

Longitudinal traces of all quality control parameters are accessible for each instrument, facilitating assessment of variability changes over time for each laboratory and instrument type. As example, we can follow the evolution of the number of peptides identified in an instrument over four years of quality control standards (Figure 2A). It can be observed that despite the existence of some spikes related to occasional malfunctions, in terms of peptide identifications the performance of Orbitrap Fusion Lumos 1 has been fairly constant over the four years for all the different fragmentation methods tested in all sites (Figure 2B). Several sites exhibited a similar number of identified peptides among them, while other sites had also a stable performance, but with lower number of identifications. This observation isn’t inherently negative, as various reasons could account for these differences, including different LC-MS setups, column types, etc. Importantly, these differences triggered discussions within the Core for Life Proteomics Working group, to identify cases where observed divergences could be explained by an under-performing instrumentation, or misconfiguration, or other examples where, on the contrary, these were the result of a conscious parametrization or setup, aimed at a specific application. These discussions made it possible to establish an open frame for information flow that resulted in an overall reduction of the coefficient of variation of the studied parameter in all instruments during a period of the harmonization study (Figure 2C, Table 1). Many of the observed differences were assigned to slightly different setups, especially in the liquid chromatographic part in which the preference of each laboratory for different column types and materials (e.g. commercial or self-made columns) as well as the inclusion or not of a pre-column justified most of the observed divergences among laboratories. Note that there are different operation modes of instrument access in the sites participating in the harmonization study, with some instruments being supervised by one single operator and dedicated exclusively to certain projects, while others are open to multiple users with different projects, including personnel in training. It was a general observation that instruments operated by multiple users and/or dedicated to multiple applications (changing setups) required more care and more frequent interventions compared to those operated by a single operator and dedicated exclusively to one type of analysis.

**Figure 2:**
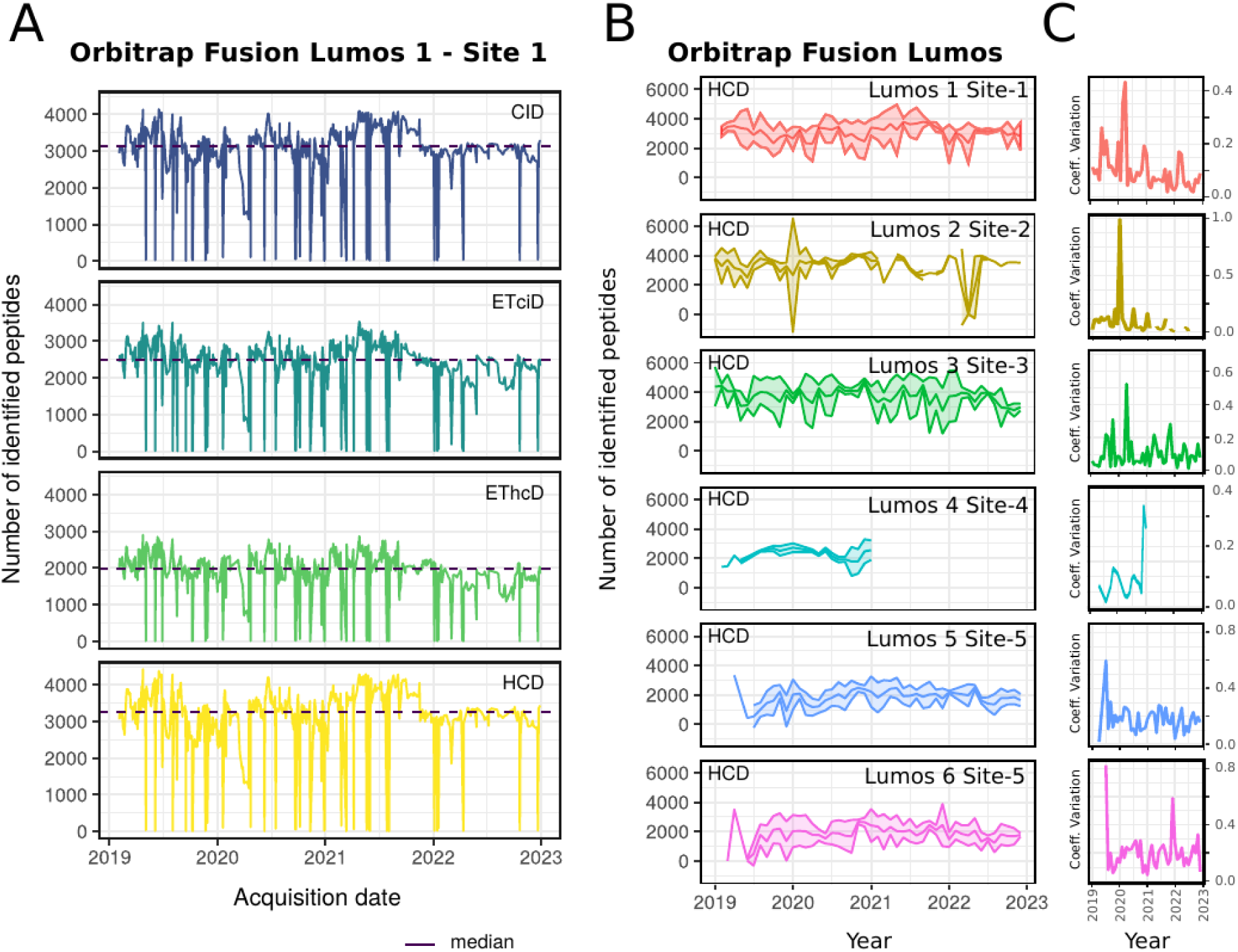
Longitudinal traces corresponding to the evolution of the number of peptides identified in an Orbitrap Fusion Lumos 1 (Site 1) instrument over four years of quality control standards in different fragmentation strategies. B) Longitudinal evolution of monthly mean and standard deviation of the number of peptides identified (HCD) in all the Orbitrap Fusion Lumos instruments monitored in the harmonization study. Points without valid values (NA, or zero) have been removed from the plot. C) Longitudinal evolution of the coefficient of variation of the number of peptides identified (HCD) in all instruments during a period of the harmonization study. Outlier points (<500 identified peptides) and points without valid values (NA, or zero) have been removed from the plot.

**Table 1:** Average number of identified peptides with HCD fragmentation in the different instruments enrolled in the harmonization study. Average and dispersion measures refer to the active period of each instrument within the harmonization study. Instruments that showed a better coefficient of variation within the last year are highlighted in green.

| Instrument | Average | Std Deviation | CV | Last year CV* | n |
| --- | --- | --- | --- | --- | --- |
| Orbitrap Eclipse 1 Site-1 | 3240.75 | 738.98 | 0.23 | 0.15 | 383 |
| Orbitrap Eclipse 2 Site-1 | 3620.26 | 566.03 | 0.16 | 0.13 | 185 |
| Orbitrap Exploris 480 1 Site-3 | 6996.19 | 2256.20 | 0.32 | 0.38 | 219 |
| Orbitrap Exploris 480 2 Site-3 | 5926.85 | 2046.72 | 0.35 | 0.35 | 75 |
| Orbitrap Exploris 480 3 Site-8 | 6202.20 | 2822.66 | 0.46 | 0.22 | 5 |
| Orbitrap Exploris 480 4 Site-4 | 5431.01 | 759.92 | 0.14 | 0.14 | 80 |
| Orbitrap Exploris 480 5 Site-4 | 4303.65 | 648.02 | 0.15 | 0.15 | 107 |
| Orbitrap Exploris 480 6 Site-4 | 5046.60 | 976.65 | 0.19 | 0.19 | 108 |
| Orbitrap Exploris 480 7 Site-4 | 5423.71 | 604.76 | 0.11 | 0.11 | 17 |
| Orbitrap Fusion Lumos 1 Site-1 | 3287.64 | 550.17 | 0.17 | 0.08 | 584 |
| Orbitrap Fusion Lumos 2 Site-2 | 3504.55 | 760.76 | 0.22 | 0.07 | 379 |
| Orbitrap Fusion Lumos 3 Site-3 | 3960.60 | 649.01 | 0.16 | 0.17 | 413 |
| Orbitrap Fusion Lumos 4 Site-4 | 2254.53 | 390.07 | 0.17 | 0.26 | 76 |
| Orbitrap Fusion Lumos 5 Site-5 | 2069.05 | 521.50 | 0.25 | 0.21 | 606 |
| Orbitrap Fusion Lumos 6 Site-5 | 2162.46 | 579.11 | 0.27 | 0.24 | 710 |
| Q Exactive HF 1 Site-3 | 9164.74 | 3062.66 | 0.33 | 0.31 | 387 |
| Q Exactive HF 2 Site-6 | 4324.44 | 1538.92 | 0.36 | 0.28 | 198 |
| Q Exactive HF 3 Site-6 | 1731.09 | 932.70 | 0.54 | NA | 11 |
| Q Exactive HF 4 Site-6 | 3576.05 | 1334.00 | 0.37 | 0.22 | 311 |
| Q Exactive HF 5 Site-6 | 2331.61 | 1044.71 | 0.45 | 0.43 | 28 |
| Q Exactive HF 6 Site-7 | 8516.96 | 1984.89 | 0.23 | 0.14 | 259 |
| Q Exactive HF 7 Site-5 | 3986.79 | 1855.96 | 0.47 | 0.43 | 793 |
| Q Exactive HF 8 Site-5 | 4185.25 | 2113.90 | 0.51 | 0.52 | 808 |
| Q Exactive HF 9 Site-5 | 4970.25 | 2250.16 | 0.45 | 0.42 | 1100 |
| Q Exactive HF-X 1 Site-8 | 10928.40 | 2550.31 | 0.23 | 0.15 | 134 |
| Q Exactive HF-X 2 Site-3 | 12374.51 | 4319.90 | 0.35 | NA | 146 |
| Q Exactive HF-X 3 Site-4 | 6821.96 | 2042.07 | 0.30 | 0.25 | 26 |
| Q Exactive HF-X 4 Site-4 | 5768.33 | 1733.71 | 0.30 | 0.11 | 132 |
| Q Exactive HF-X 5 Site-4 | 5753.05 | 2016.13 | 0.35 | 0.05 | 63 |
| Q Exactive HF-X 6 Site-4 | 6178.26 | 2209.59 | 0.36 | 0.08 | 65 |

\* CV of the last year of activity for each given instrument

#### Causation of performance drift

Despite the global tendencies, we also focused on periods of unexpected under-performance. During the course of this harmonization study, it became clear that the quality control data collection and monitoring on the QCloud server improved the diagnosis of LC-MS system technical unconformities within the participant laboratories. Indeed, many of the problems in the LC-MS systems were detected in the liquid chromatographic part. One particular case where this benefit was perceived was on the daily operation of two separate LC flow paths coupled to the same mass spectrometer (Orbitrap Q Exactive HF) through a dual column nanospray source system setup on Site 5. LC-MS performance metrics on both flow paths were expected to correlate, given the shared MS instrument and the use of the same acquisition method, a common QC sample, and identical buffers and autosampler. This close association between both flow paths can indeed be perceived when looking at a longitudinal plot of peptide identification rates for the whole course of the study (Figure 3A). In an example close-up period from early 2020 (Figure 3A, left inset), data points from flow paths A and B are seen to go hand in hand most of the time. A histogram of the difference in the same QC metric (number of identified peptides) between these two separate LC flow paths is nicely centered around zero, indicating an overall equal performance of both paths (Figure 3B). Nonetheless, deviations in peptide identifications between the two flow paths could go as high as 6,209 (i.e. ∼150 % of the MS instrument’s mean value (including lab system A and B) of 4,147). By continuously monitoring the relative performance of both flow paths, LC-related problems can be quickly detected once QC metrics start to diverge between the two. As an example of the latter, we focus on a period of five weeks from early 2021 (Figure 3A, right inset). Between January 8 and 18, peptide identification rates were consistently lower for flow path B. This behavior steered an LC intervention on this flow path and indeed, after the installation of a new column on January 18 a major improvement on peptide ID rates was observed (a 109 % increase from 2,760 to 5,790). After this intervention, the performance also got worse for flow path A. Again, a change of analytical column at the beginning of February solved the situation, as detected by a 72 % increase (from 3,458 to 5,980) in peptide ID rates on the following QC.

**Figure 3:**
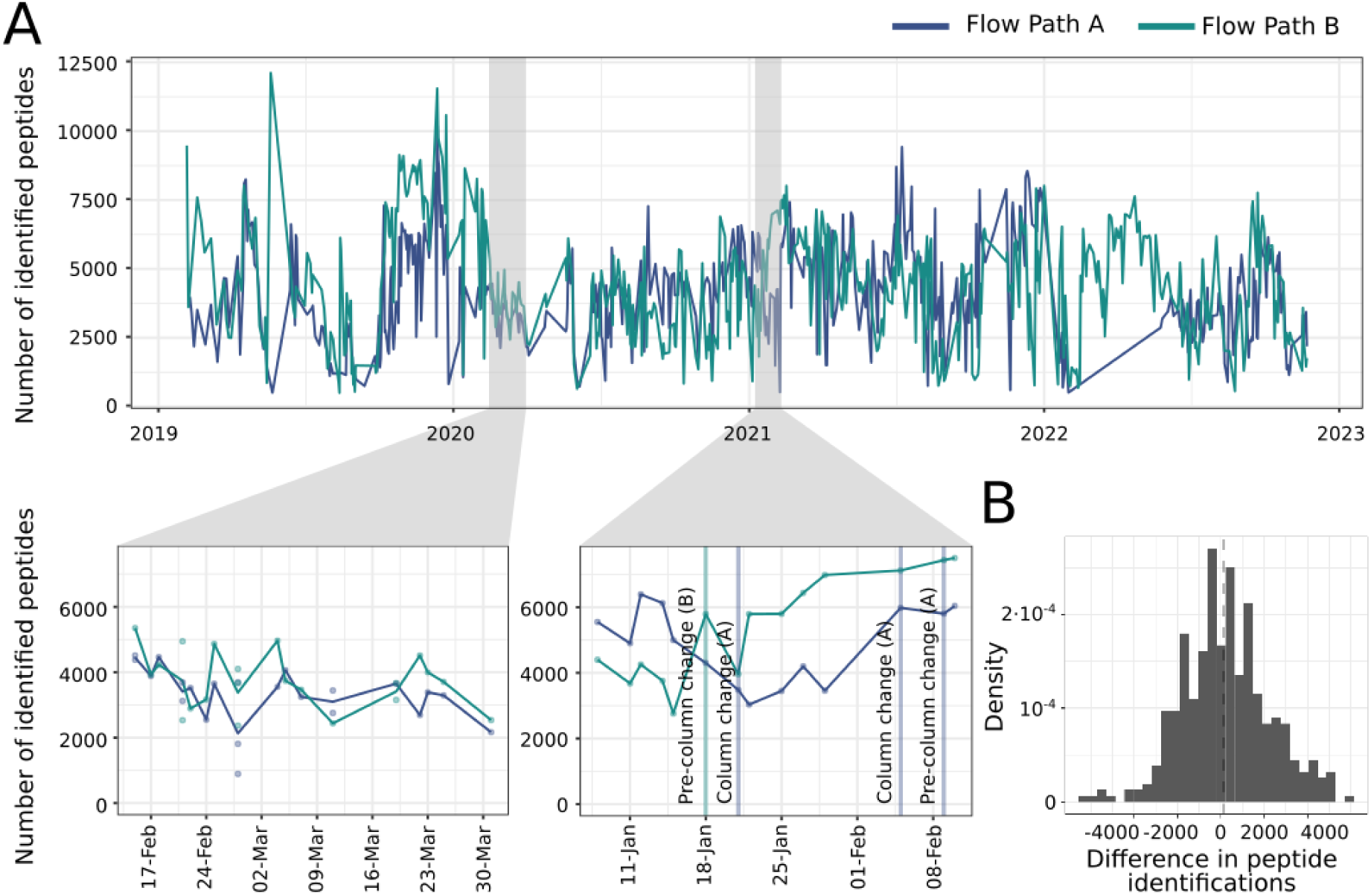
Case study on a dual column instrument set up. (A) Longitudinal measurements of peptide identification rates in two lab systems connected to the same Orbitrap Q Exactive HF instrument. Top panel. Full period of harmonization. Left inset: Uneventful representative short period from 14^th^ Feb 2020 to 31^st^ March 2020. Right inset: Short period with non-conformities followed by key technical interventions: 2021/01/08-2021/02/10. (B) Histogram of the difference between peptide identification rates in flow path A and B, during the course of the study.

Another example was reported by Sites 1 and 6 in which QCloud indicated a low number of identified peptides and proteins but with normal total and extracted ion chromatogram MS1 areas of many different peptides. Once the gas pressure used for peptide fragmentation was checked as correct, the report pointed to a problem with an electronic instrument control board responsible for peptide fragmentation. This triggered a call to the service engineer, who certified the problem and replaced the board.

Finally, sometimes differences in performance were due to unintentional changes in the parameters of the method (e.g. Site 3, Orbitrap Fusion Lumos 3) or even more subtle aspects like an upgrade of the Tune software version of an instrument, resulting in unexpected behavior even when using the exact same data acquisition method. An example of the latter was documented in the Orbitrap Fusion Lumos 2 from Site 2 in spring 2019 when comparing it to the non-upgraded tune version of the same instrument type at Site 1 (upgrade from Tune version 3.0 to 3.1).

The combination of a close monitoring of the QCloud system with information exchange among participants helped each site to detect problems in calibration, ion optics, spray stability, hardware, or chromatographic alterations. During the execution of the harmonization study, other quality control parameters, not explicitly evaluated in the study, were also suggested as valuable factors to expand the range of diagnostic tools in LC-MS systems within our working group. Examples of these high-level parameters are the ratio among different ion charges, automatic detection of main contaminants, the proportion of peptides identified in the hydrophilic, middle, and hydrophobic parts of the gradient, or the monitoring of the real resolution on the instrument, among others.

While it is beyond the scope of this manuscript to describe the myriad problems and extensive discussions encountered throughout our study, it is pivotal to highlight the immense benefits collected from this harmonization study. The collaborative sharing of reports and quality control values not only facilitated the identification of the causes behind the under performance of our systems, but also made us aware of the impact that certain decisions on LC-MS setup had in instrument performance. Among others, they triggered changes in chromatography setups, and helped us identify and document persistent problems in certain instruments. Overall, the harmonization study has allowed us to significantly reduce the variability inherent to our LC-MS systems, and the collective discussion of the encountered problems and divergences has given us greater confidence in accurately identifying LC-MS related problems. Consequently, this allows us to reach out to service engineers more reliably for targeted interventions, as opposed to scheduled regular interventions, which, in the context of an already optimally functioning system, may only minimally impact performance.

### Establishment of Community Reference Value

Finally, the number of the identified peptides obtained by the same type of instrument were compared (Figure 4), and based on the longitudinal data from the different instruments that participated in the harmonization study, we established an average reference value and a range for each of the 109 quality control parameters monitored in this study per instrument type. These reference values are formed by the average, the median, the standard deviation, the coefficient of variation (%), and the quantile values 25, 50 and 75% for each parameter in each type of instrument (Supplementary Table ST3).

**Figure 4:**
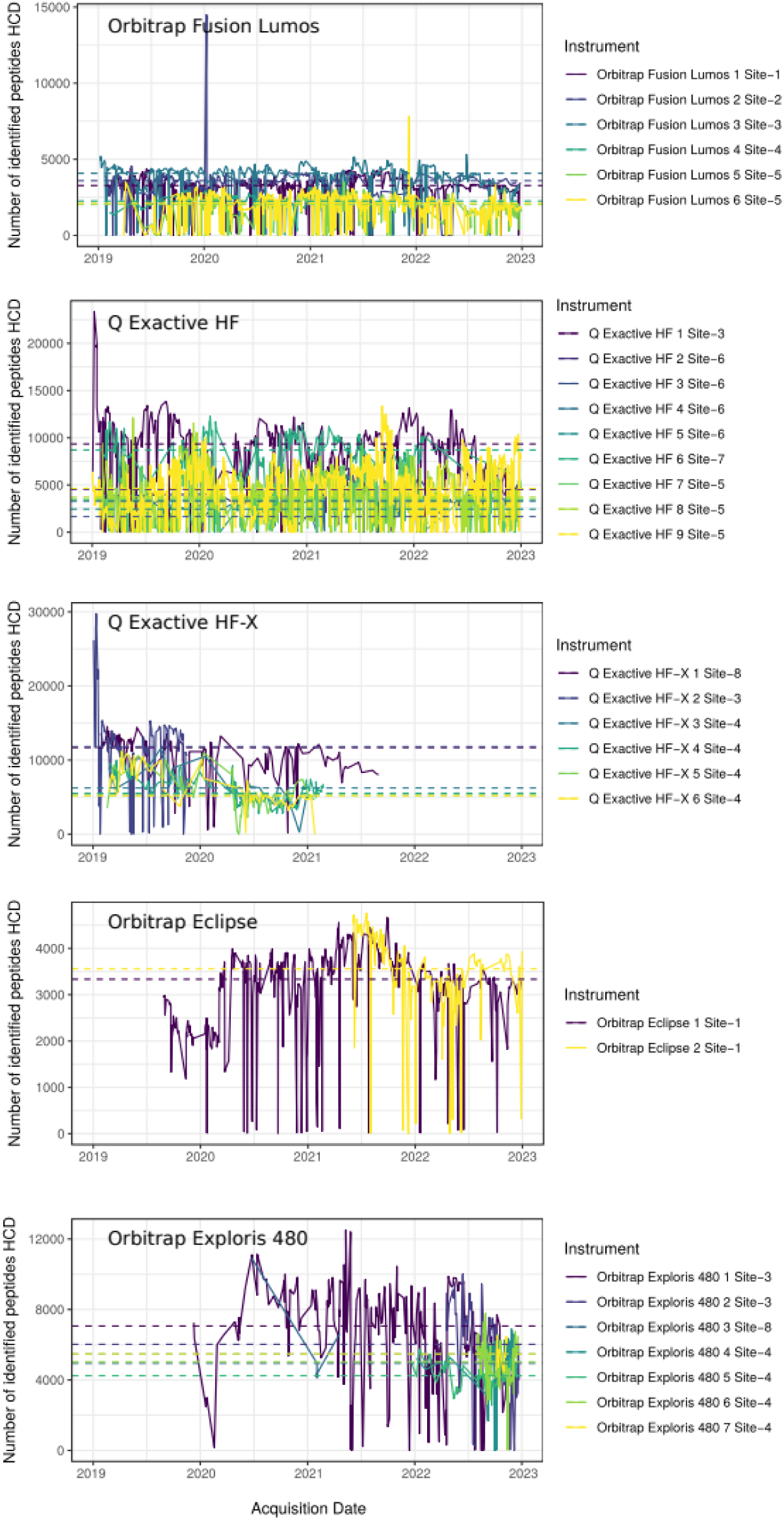
Longitudinal comparison of the number of identified peptides (HCD fragmentation obtained with the same type of instruments that participated in the harmonization study. The dotted line represents the median for the individual instruments.

The reference values for the Orbitrap Fusion Lumos, Orbitrap Q Exactive HF, and Orbitrap Q Exactive HF-X are based on an extensive time period spanning from the beginning of 2019 to the end 2022 in which the harmonization study has been conducted. In contrast, for newer instrument models, such as the Orbitrap Eclipse and the Orbitrap Exploris 480, the reference values have been drawn from shorter time series and fewer units since they were incorporated later into the harmonization study as they were acquired and installed in the various Core for Life laboratories. In any case, it should be noted that the reference values obtained span a broad range of configurations found in the various Core for Life laboratories, and therefore, they are not strict threshold values as they are not the result of a standardization process, but rather the result of four years of a harmonization study. These values (median) have been incorporated into the QCloud graphical user interface as a reference value achieved in the Core for Life Proteomics Working Group, and should function as quality indicator that other laboratories can refer to when considering the same type of instrument, method, and standard control sample (Supplementary Figure S3).

All the mass spectrometry proteomics data associated to this study have been deposited at the ProteomeXChange Consortium via the PRIDE repository with identifier PXD049444 (Vizcaíno et al, 2014).

## Conclusion

In our study, we performed a coordinated effort to establish a common quality control framework within the proteomics core facilities of the Core for Life alliance. The goal was to enhance the consistency of quality control results, thereby pinpointing standard reference values for the community, fluctuations in performance within labs, potential heterogeneity causes, and areas for enhancement. As a result of our harmonization study, we established a constant exchange of information and peer-discussion around under-performing periods that facilitated the accurate identification of LC-MS related problems and eventually allowed us to reduce the variability in our LC-MS systems. Beyond the assessment of variability both within and between labs, we report a new quality control standard for monitoring the performance of nano liquid chromatography-mass spectrometry (LC-MS) in proteomics applications, a collection of community reference values, and a 4-year longitudinal dataset of quality control readings from multiple instruments and laboratories, beneficial for forthcoming computational proteomics studies focused on quality control. Far from being a time-limited endeavor, this initiative established a mode of operation and information exchange within the Core for Life Proteomics Working group. Consequently, it represents a sustained effort that will persist and adapt over time, incorporating new LC-MS systems in the market as they become available across different institutes.

## Abbreviations

LC-MS: Liquid chromatography coupled to mass spectrometry
HCD: Higher-energy collisional dissociation
ETD: Electron-transfer dissociation
EThcD: Hybrid electron-transfer/higher-energy collision dissociation
CID: collision-induced dissociation

## Acknowledgements

The CRG/UPF Proteomics Unit is part of the Spanish Infrastructure for Omics Technologies (ICTS OmicsTech). We acknowledge support of the Spanish Ministry of Science and Innovation through the Centro de Excelencia Severo Ochoa (CEX2020-001049-S, MCIN/AEI /10.13039/501100011033), and the Generalitat de Catalunya through the CERCA programme and The Departament de Recerca i Universitats de la Generalitat de Catalunya (2021SGR01225). Région Ile-de-France and Fondation pour la Recherche Médicale grants (to D.L.), and ANR-21-CE35-0007 (to M.M). We are grateful to all the members of the respective proteomics laboratories for technical support and useful discussion, including Dr. Henrik Thomas, and Dr. Ignacy Rzagainski (MPI-CBG), Vanessa Masson and Florent Dingli (Institut Curie), Antje Dittmann, Claudia Fortes, Peter Gehrig, Jonas Grossmann, Tobias Kockmann, Laura Kunz, Chia-wei Lin, Sibylle Pfammatter, Bernd Roschitzki, Witold E. Wolski, and Simone Wüthrich (FGCZ), Thibault Chaze and Quentin Giai Gianetto (Institut Pasteur), Gerhard Dürnberger, Richard Imre, Elisabeth Roitinger, Ines Steinmacher, Michael Schuitzbier, Susanne Opravil, Gabriele Krssakova, and Florian Stanek (IMP), Evy Timmerman, Sara Dufour, Katie Boucher, Delphi Van Haver, Jarne Pauwels (VIB), and Eva Borràs, Guadalupe Espadas, Amanda Solé, Olga Pastor, Julia Morales-Sanfrutos, Enrique Alonso, Selena Fernandez, and Marc Serret (CRG).

## Supplementary Information

**Supplementary Figure S1:**
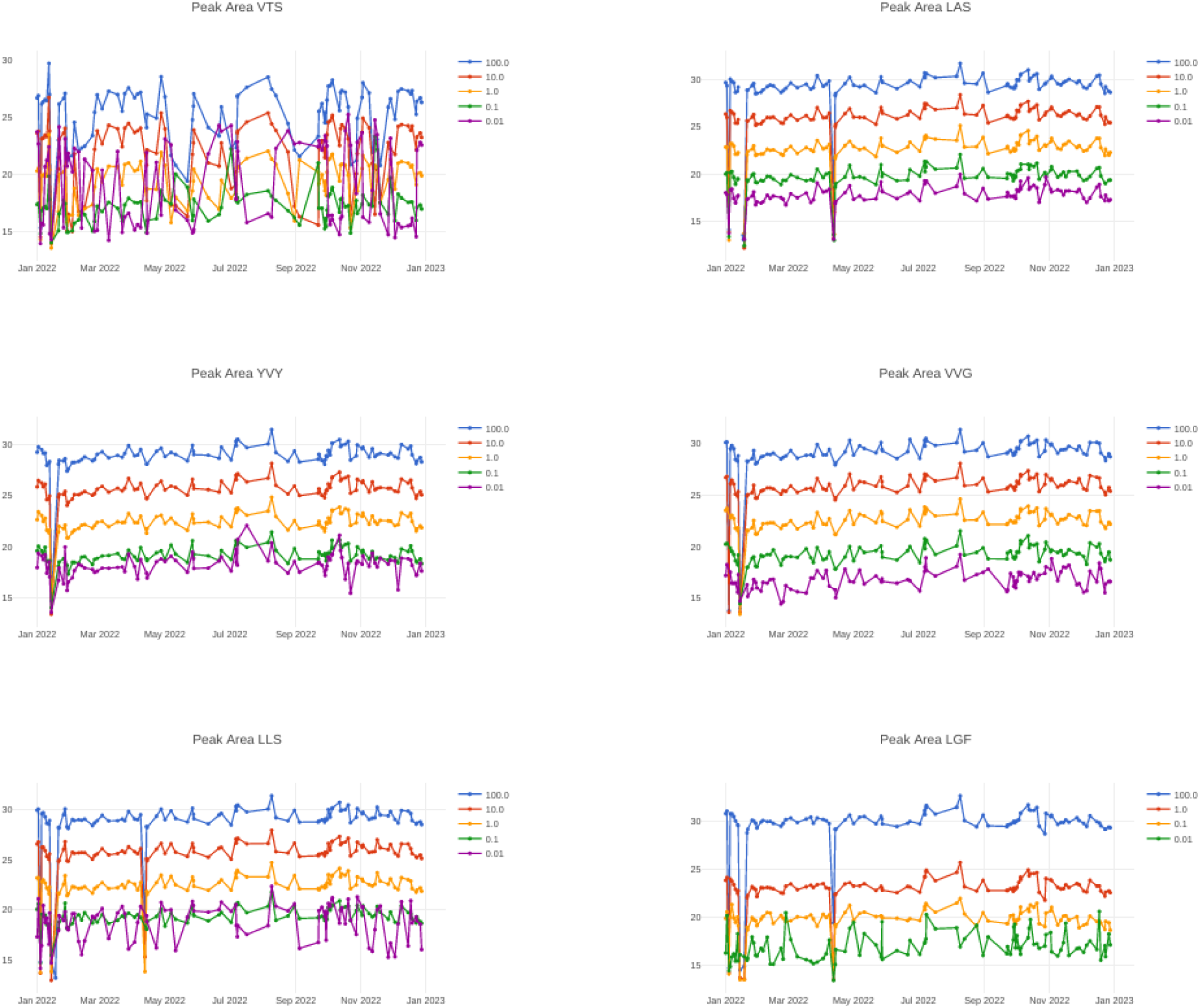
New control charts developed in QCloud to show the intensity over time of the five different isotopologue peptides included in the QC4Life Standard Mix.

**Supplementary Figure S2:**
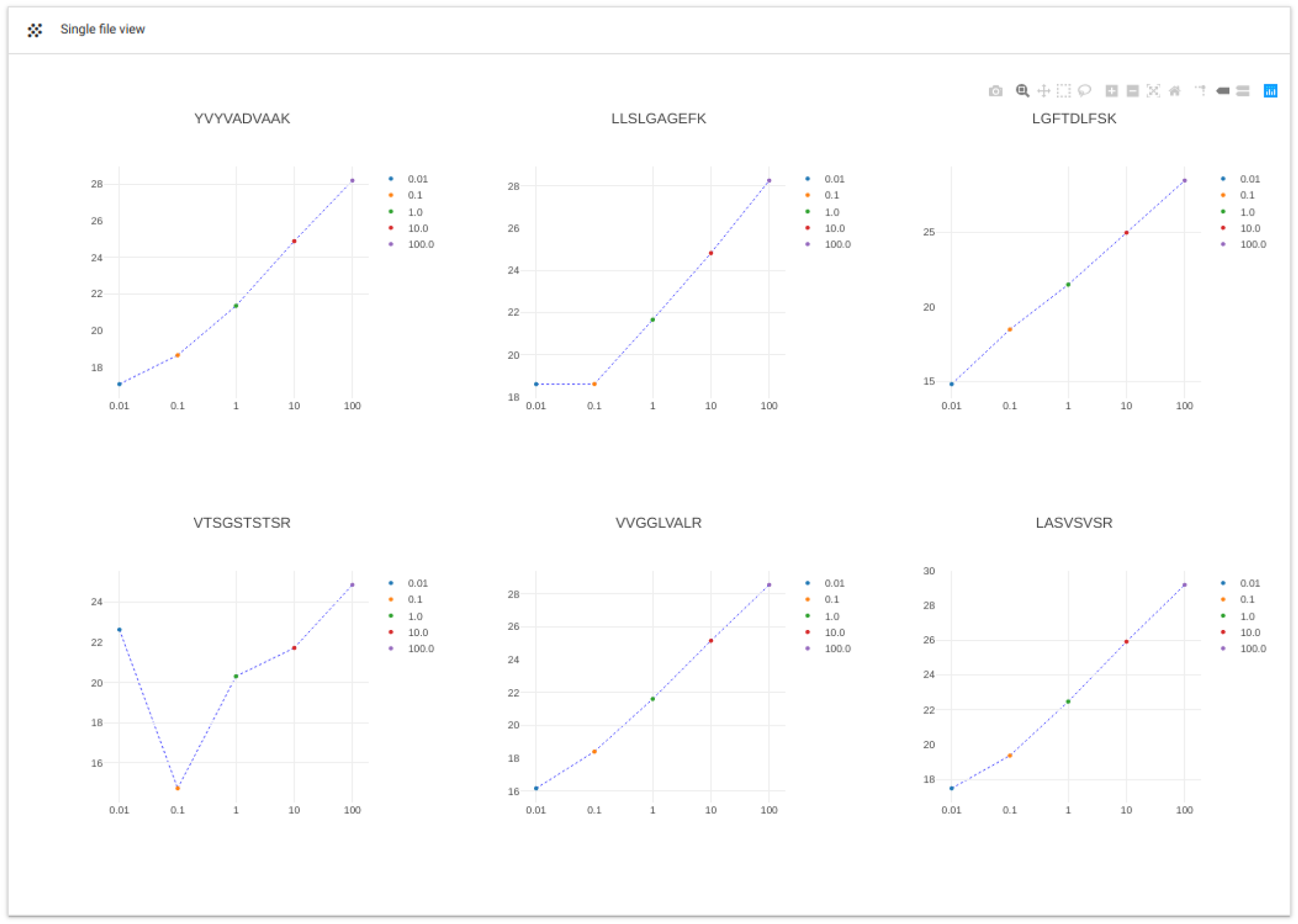
New single-file view developed in QCloud to show the intensity over time of the five different isotopologue peptides included in the QC4Life Standard Mix within each particular sample.

**Supplementary Figure S3:**
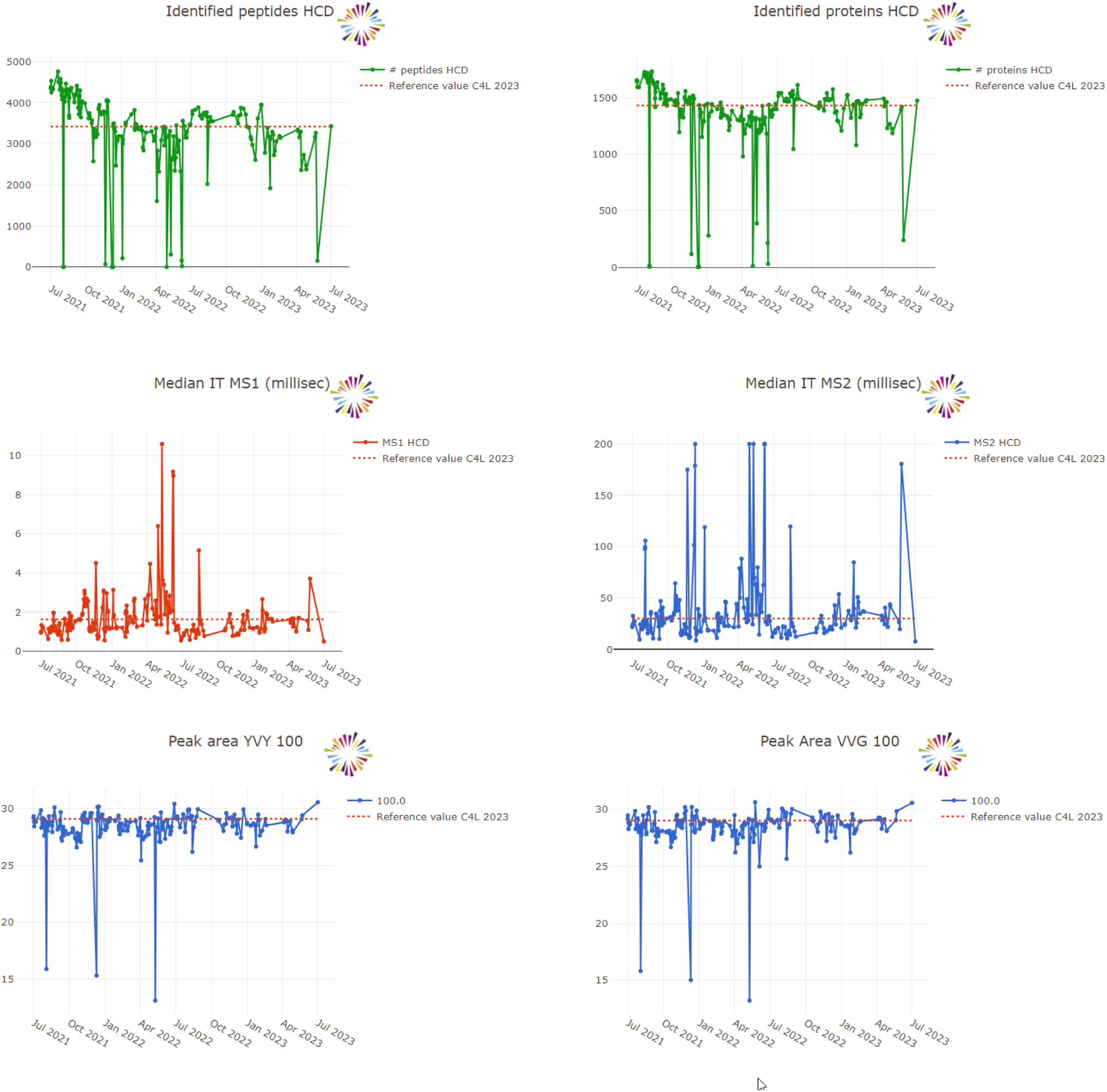
Examples of reference values from the harmonization study implemented in the QCloud graphical user interface to function as quality indicators to which other laboratories can refer when considering the same type of instrument, methods, and standard control sample.

**Report 1:** Example of a report that were automatically generated every week and shared among all the proteomics laboratories of the harmonization study within the Core for Life alliance.

**Supplementary Table ST1:** List of quality control parameters extracted during the harmonization study

**Supplementary Table ST2:** Results data with all the quality control parameter values for the individual quality control raw files analyzed during the harmnization study.

**Supplementary Table ST3:** Reference values, including central and dispersion measures for each instrument type corresponding to the 109 quality control parameters monitored in the harmonization study.

## C4L Harmonization Study - weekly report

14 May, 2020

This report is a summary of the QC data extracted from QCloud2.0 during the week of 04/05/2020 to 10/05/2020, for the QC03 analytical runs of the lab systems included in the C4L harmonization study.

It is intended for the C4L-Proteomics members, in particular those more directly involved with the generated data, to evaluate their lab systems’ performance relative to that of other nodes.

All plots show weekly median values per lab system. Error bars correspond to means ± standard deviation. For each instrument type, tables with total numbers of PSMs, peptides and proteins per analytical run are now included, next to the ‘Total numbers’ plot.

The same color code for the partner nodes will be used throughout the report.

In the peak area plots, the black regression line corresponds to the theoretical ideal line (Rsquared=1), while the blue regression line is a weighted least squares linear regression line. Peak area values from the highest abundance isotopologues were used as a reference to generate these fitted lines.

### Orbitrap Fusion Lumos

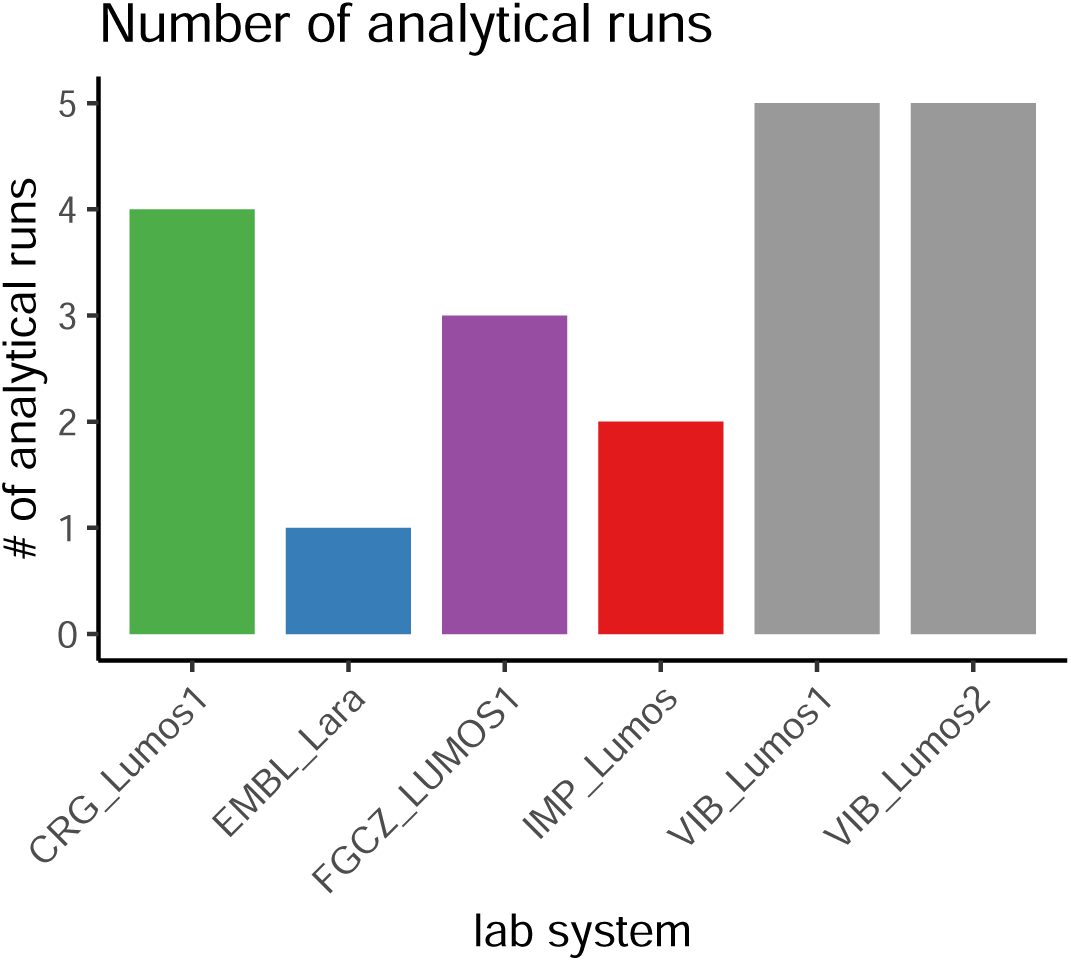

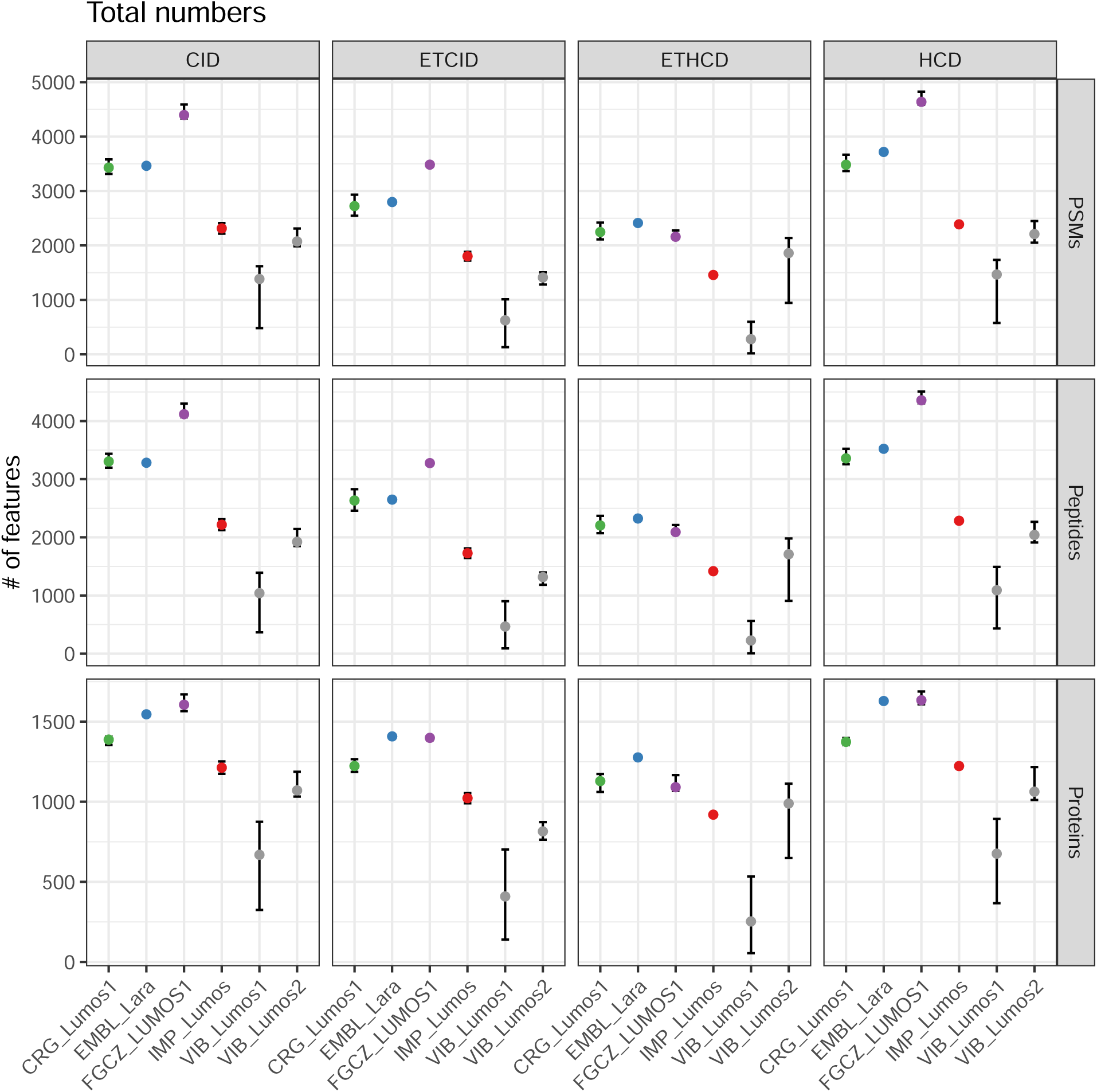

**Table 1:**
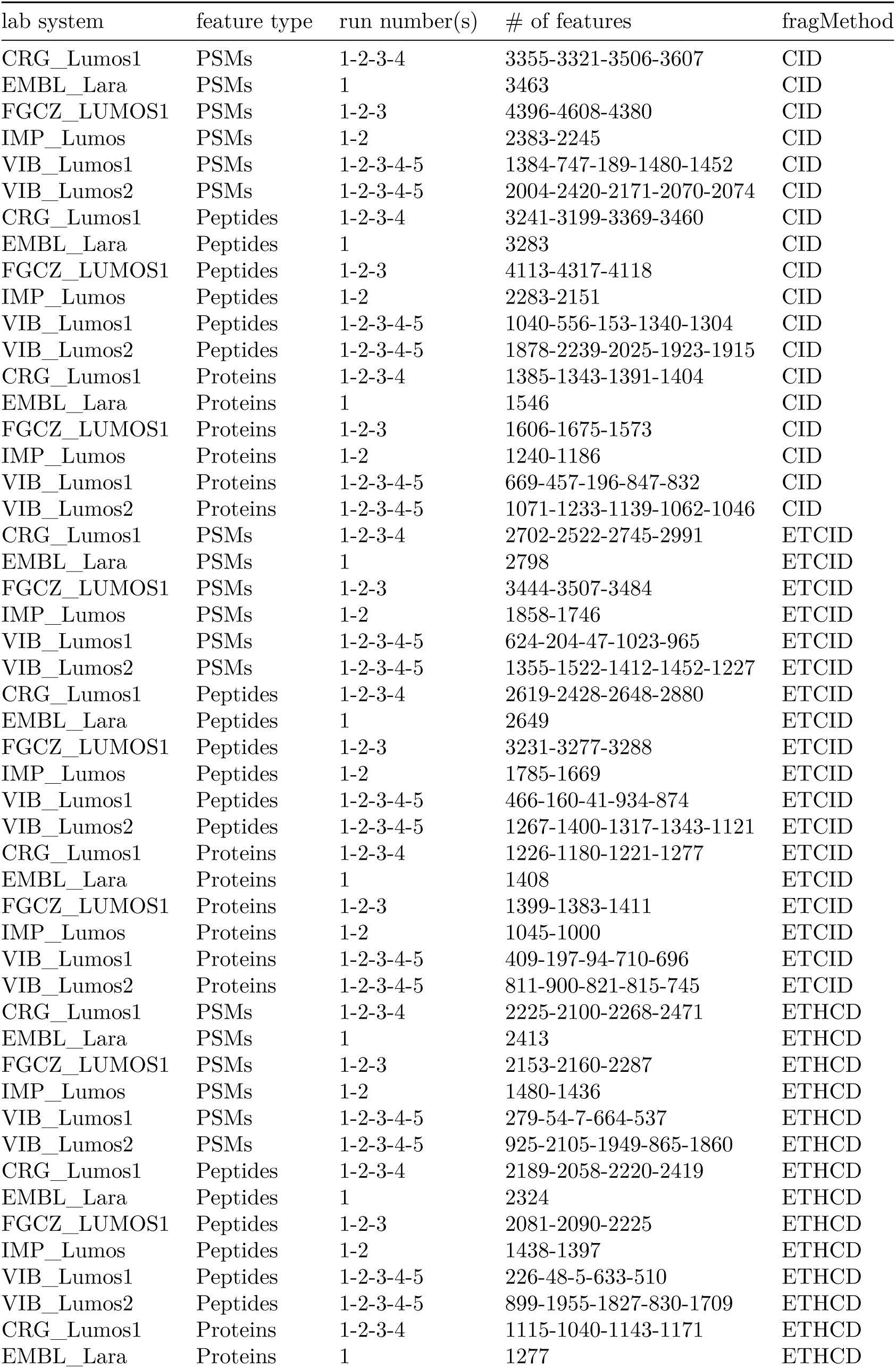
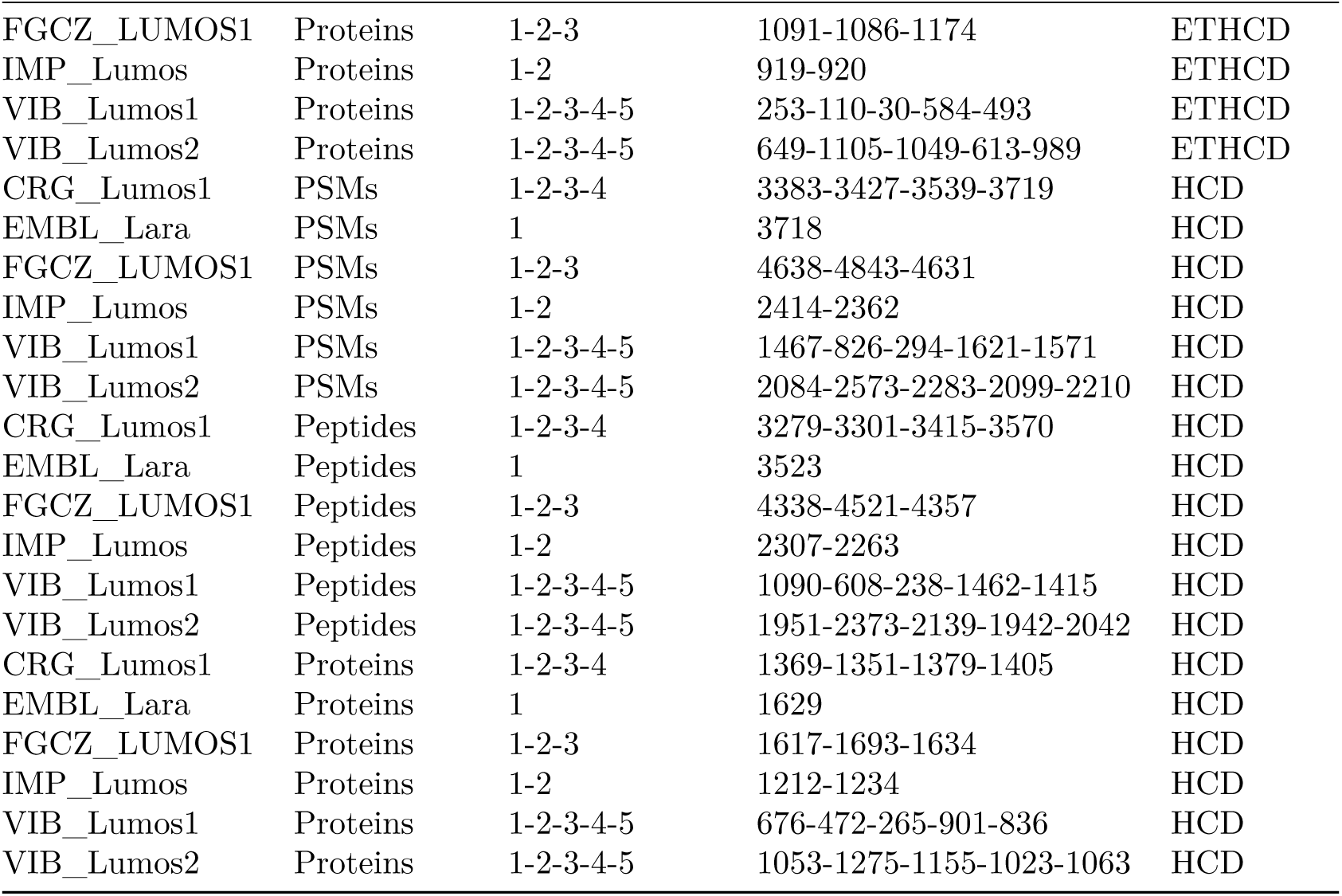
Number of features per lab system.

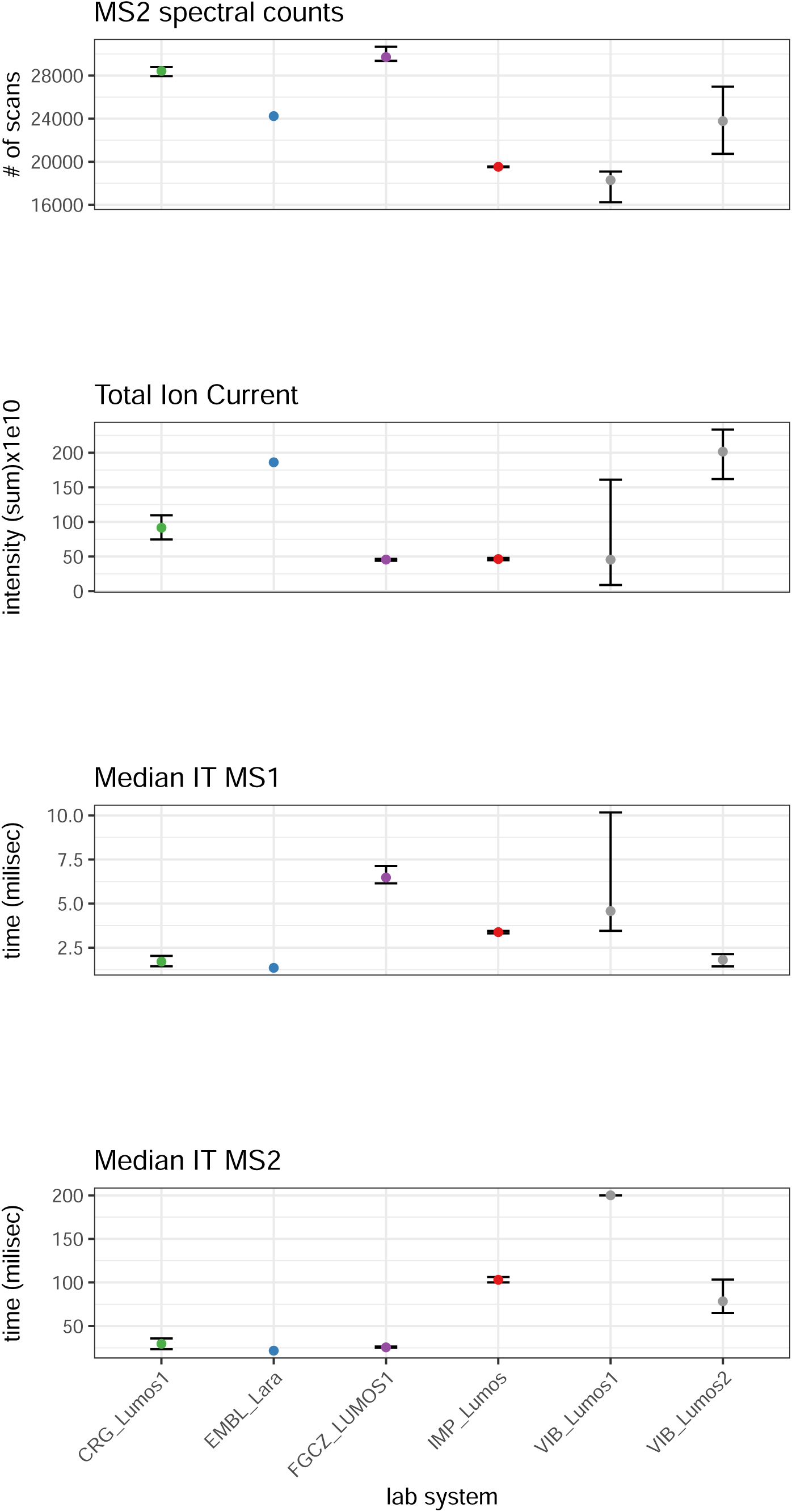

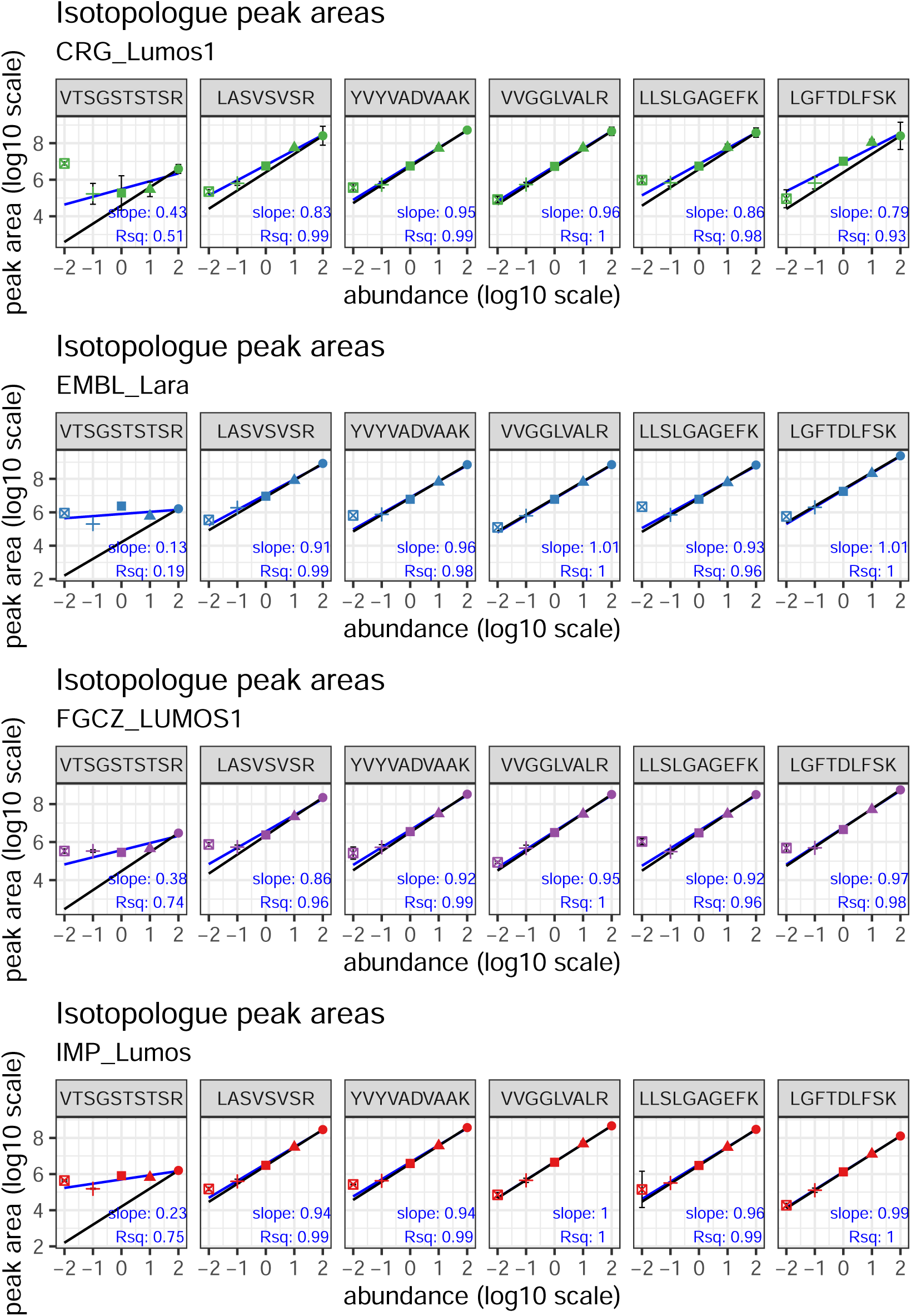

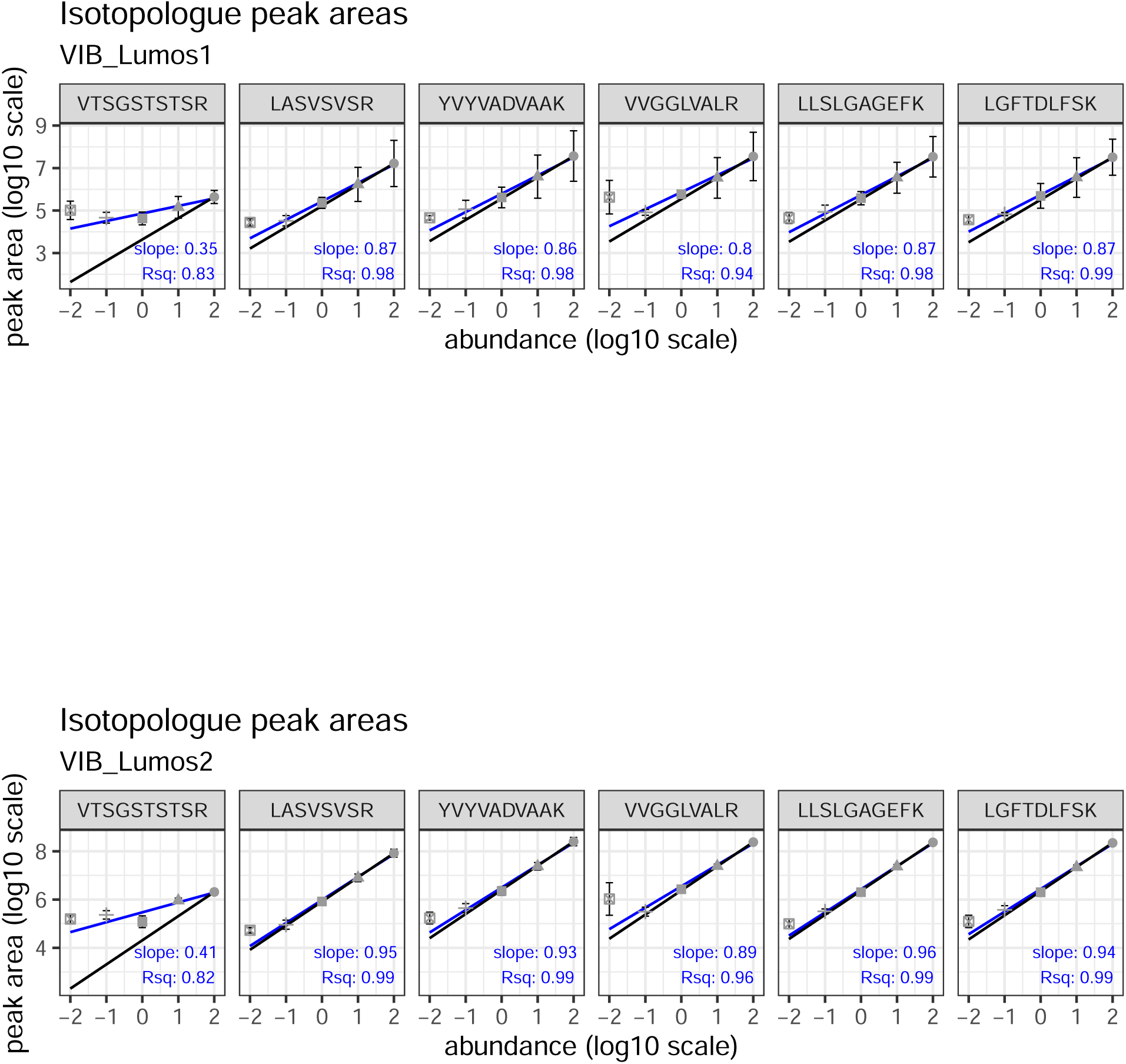

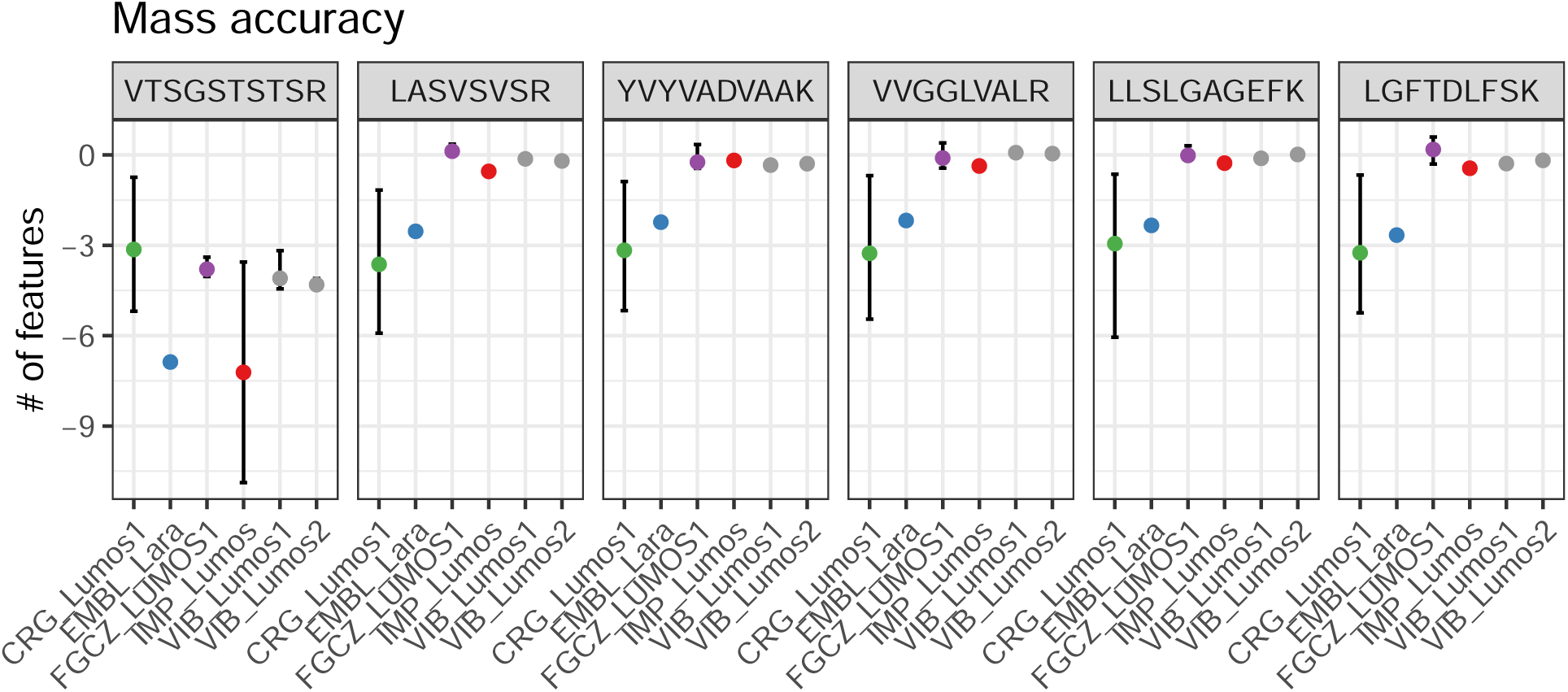

### QExactive HFX

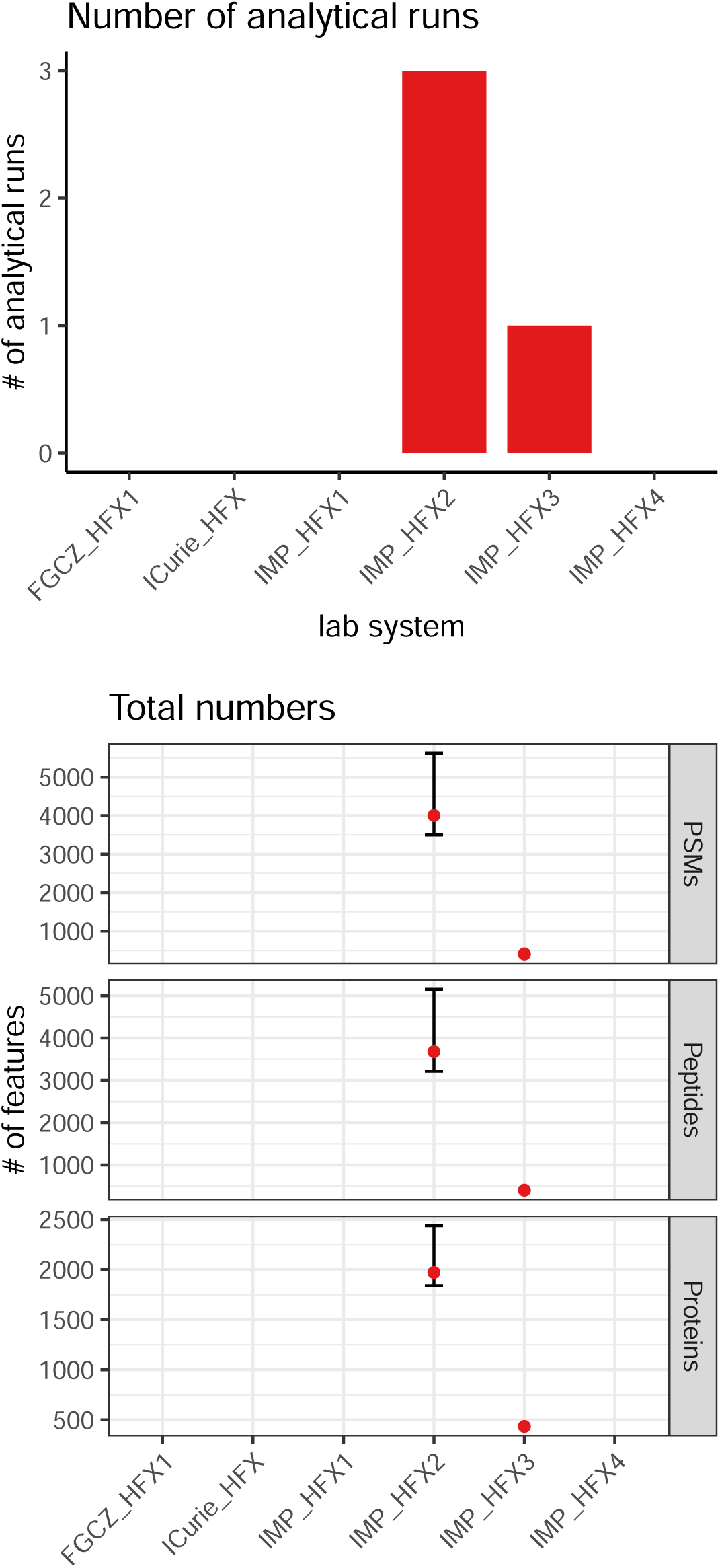

**Table 2:** Number of features per lab system.

| lab system | feature type | run number(s) | # of features |
| --- | --- | --- | --- |
| IMP_HFX2 | PSMs | 1-2-3 | 5781-4002-3891 |
| IMP_HFX3 | PSMs | 1 | 410 |
| IMP_HFX2 | Peptides | 1-2-3 | 5296-3675-3574 |
| IMP_HFX3 | Peptides | 1 | 405 |
| IMP_HFX2 | Proteins | 1-2-3 | 2485-1958-1972 |
| IMP_HFX3 | Proteins | 1 | 435 |

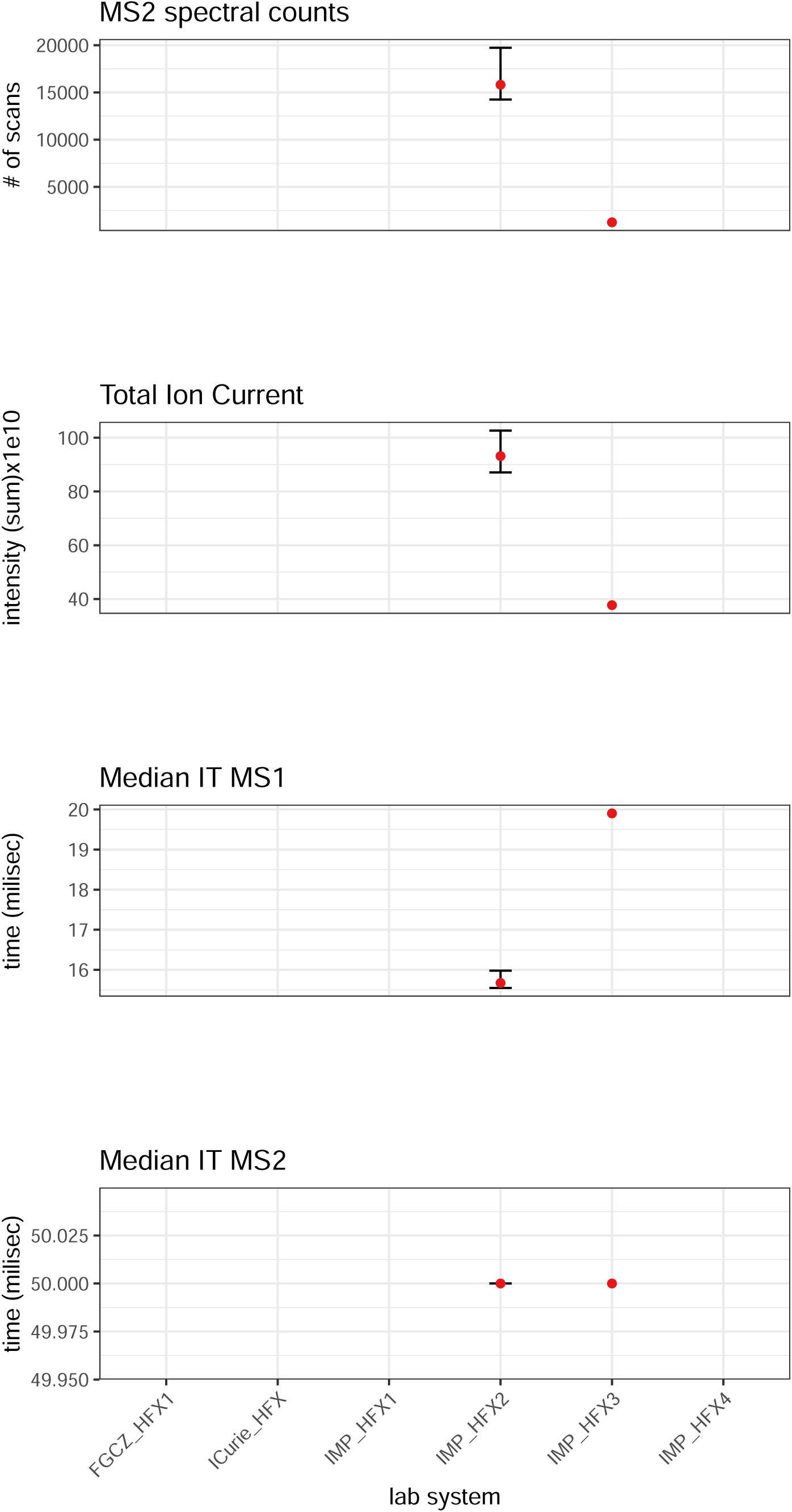

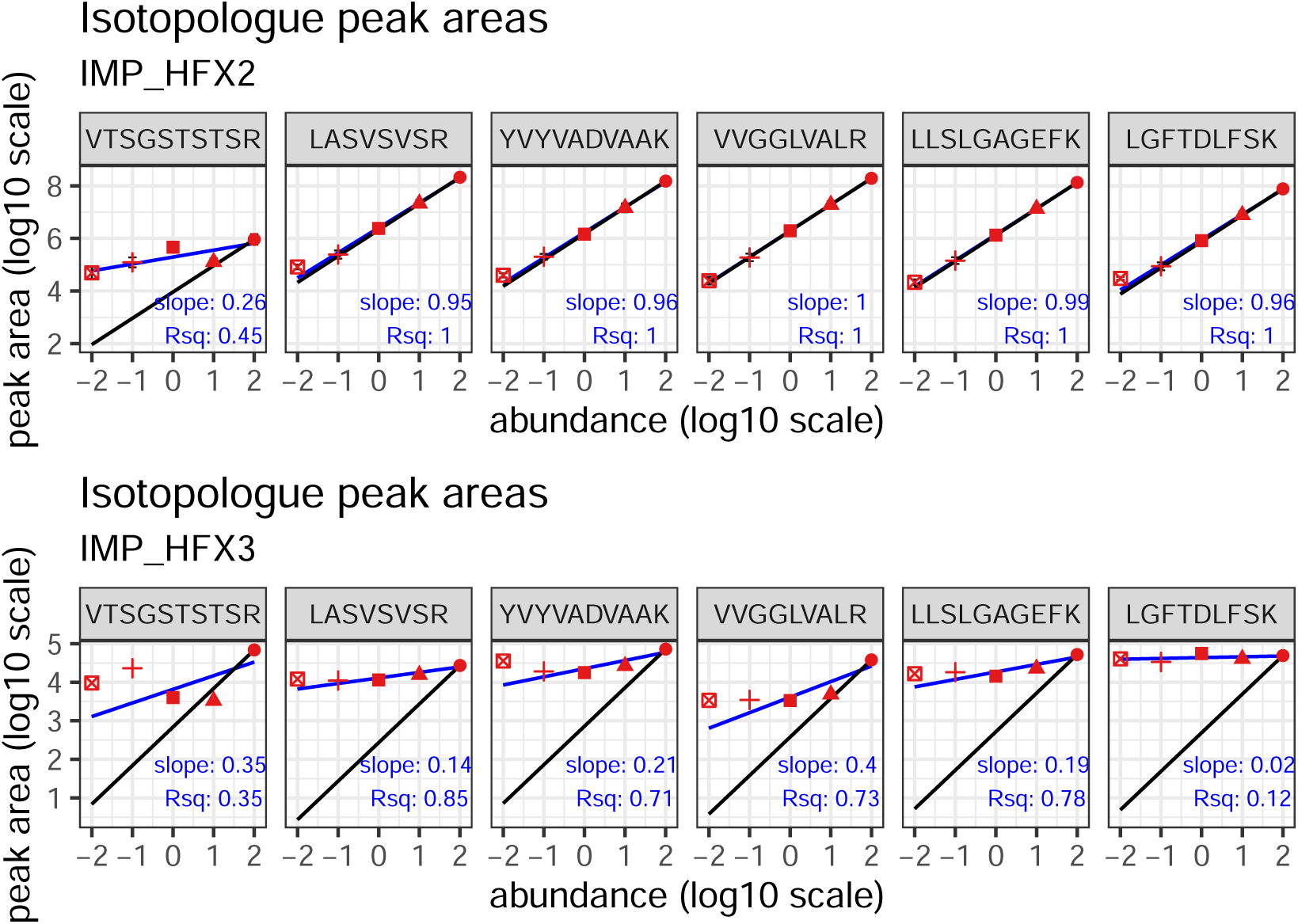

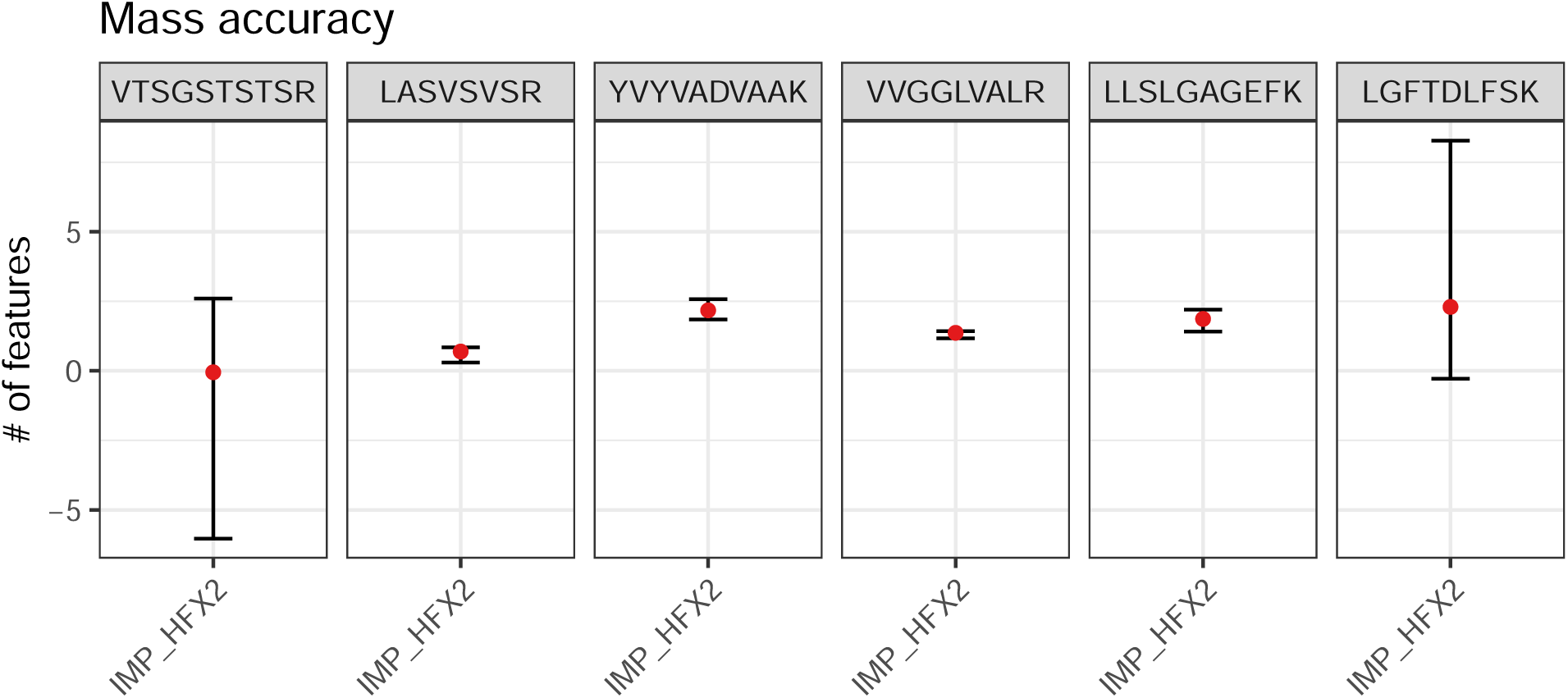

### QExactive HF

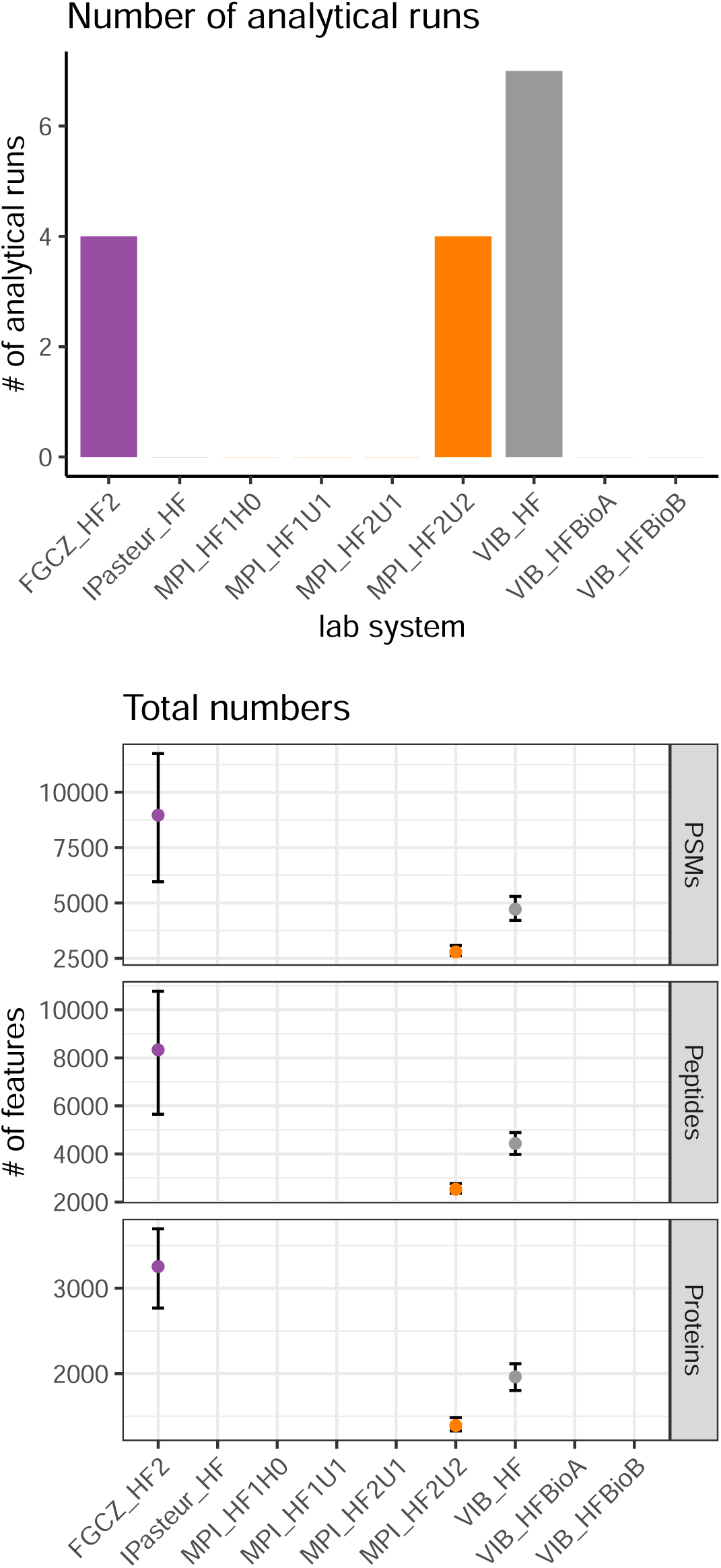

**Table 3:**
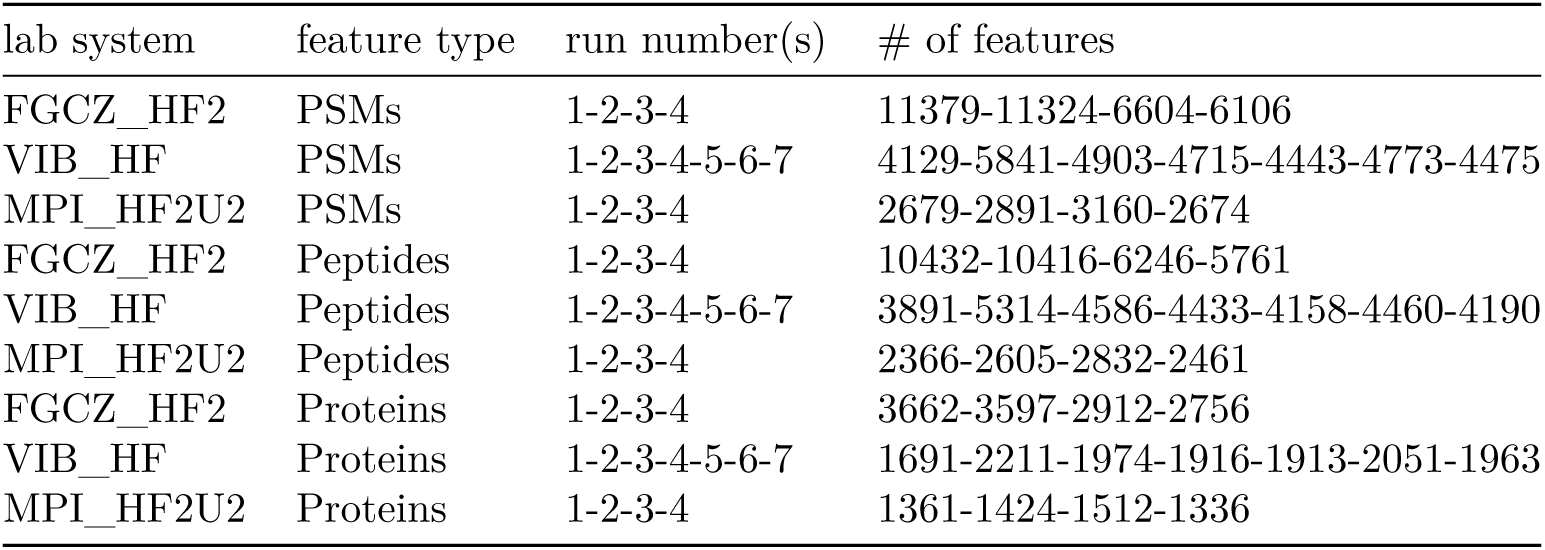
Number of features per lab system.

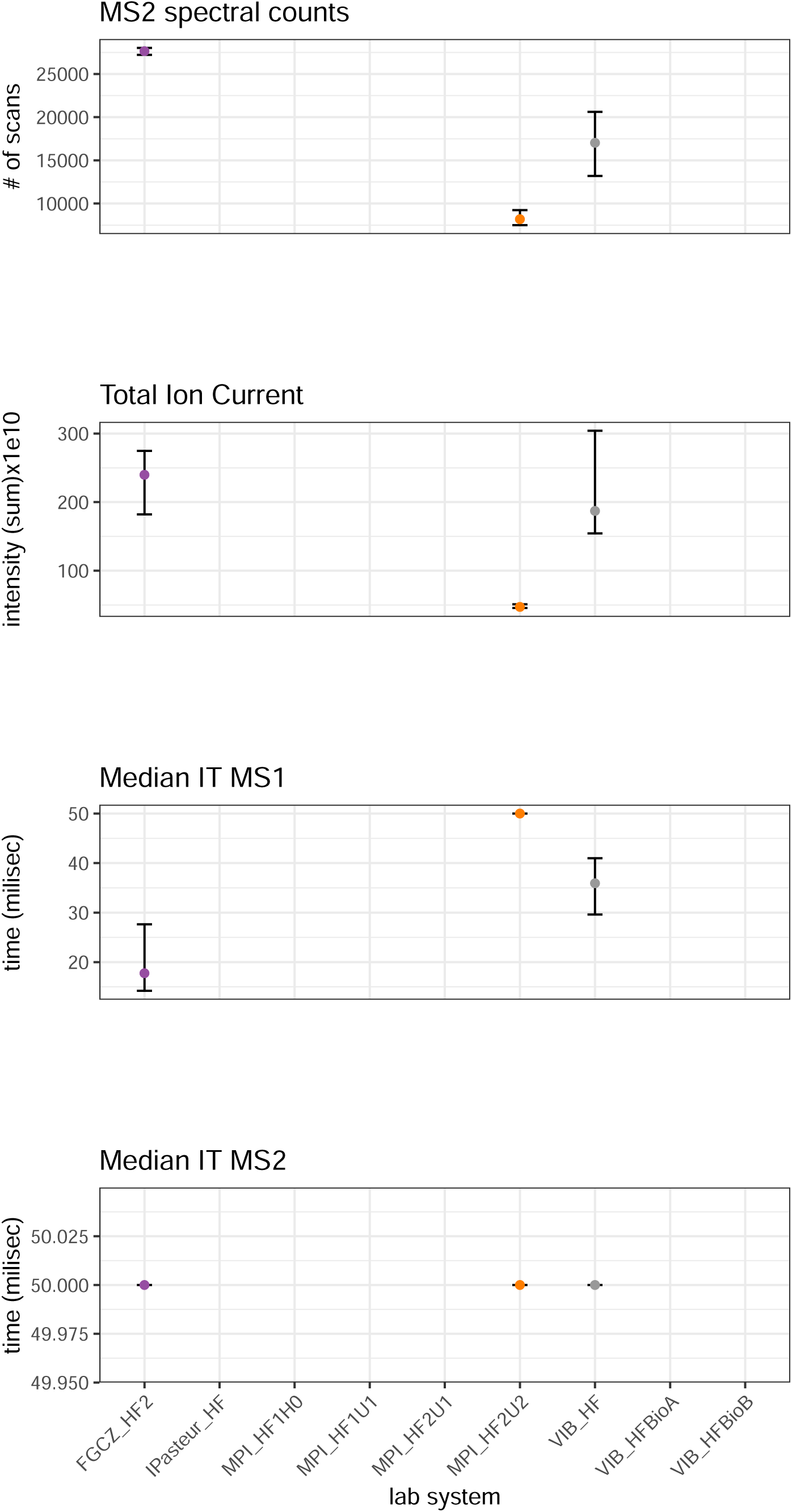

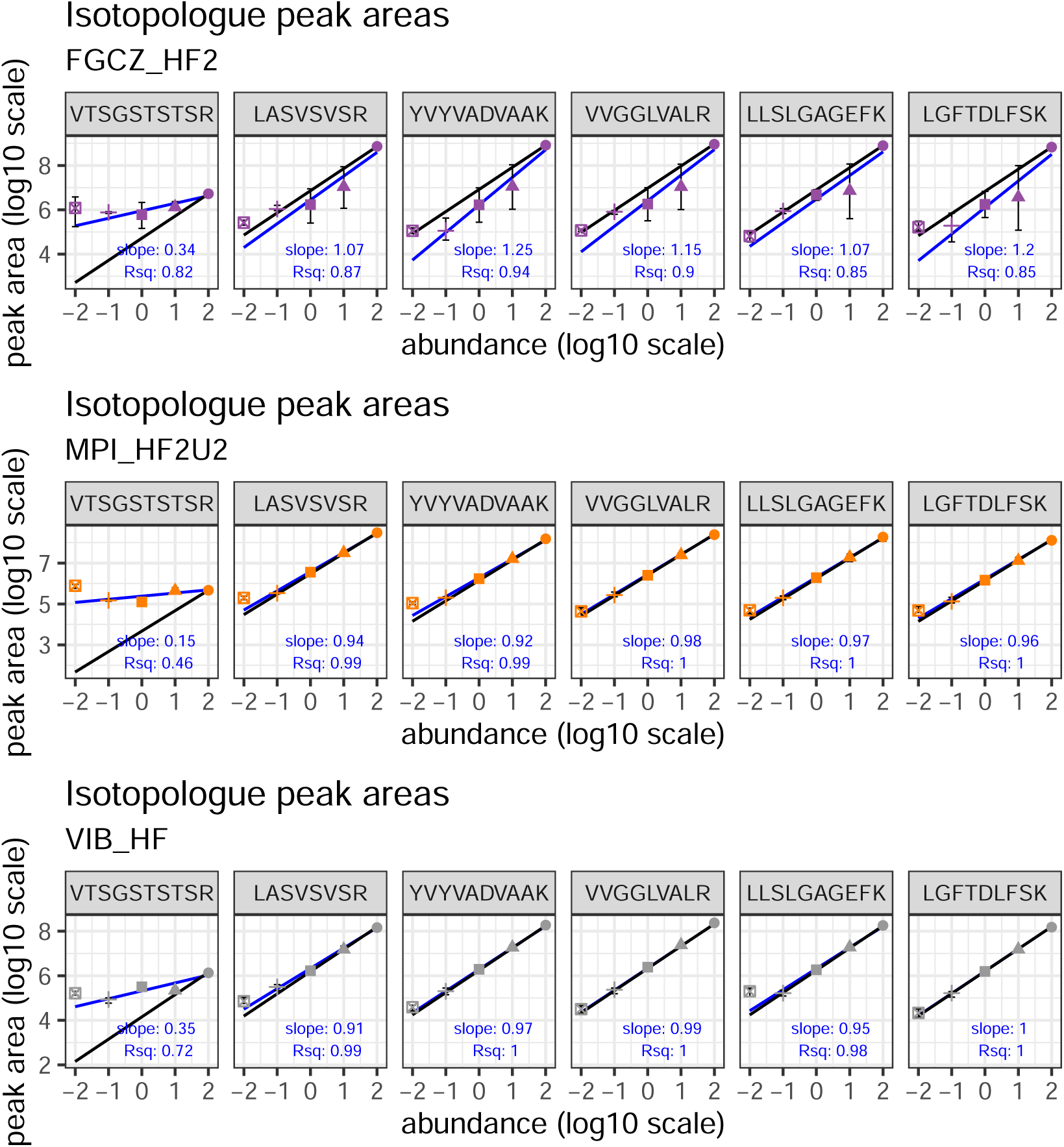

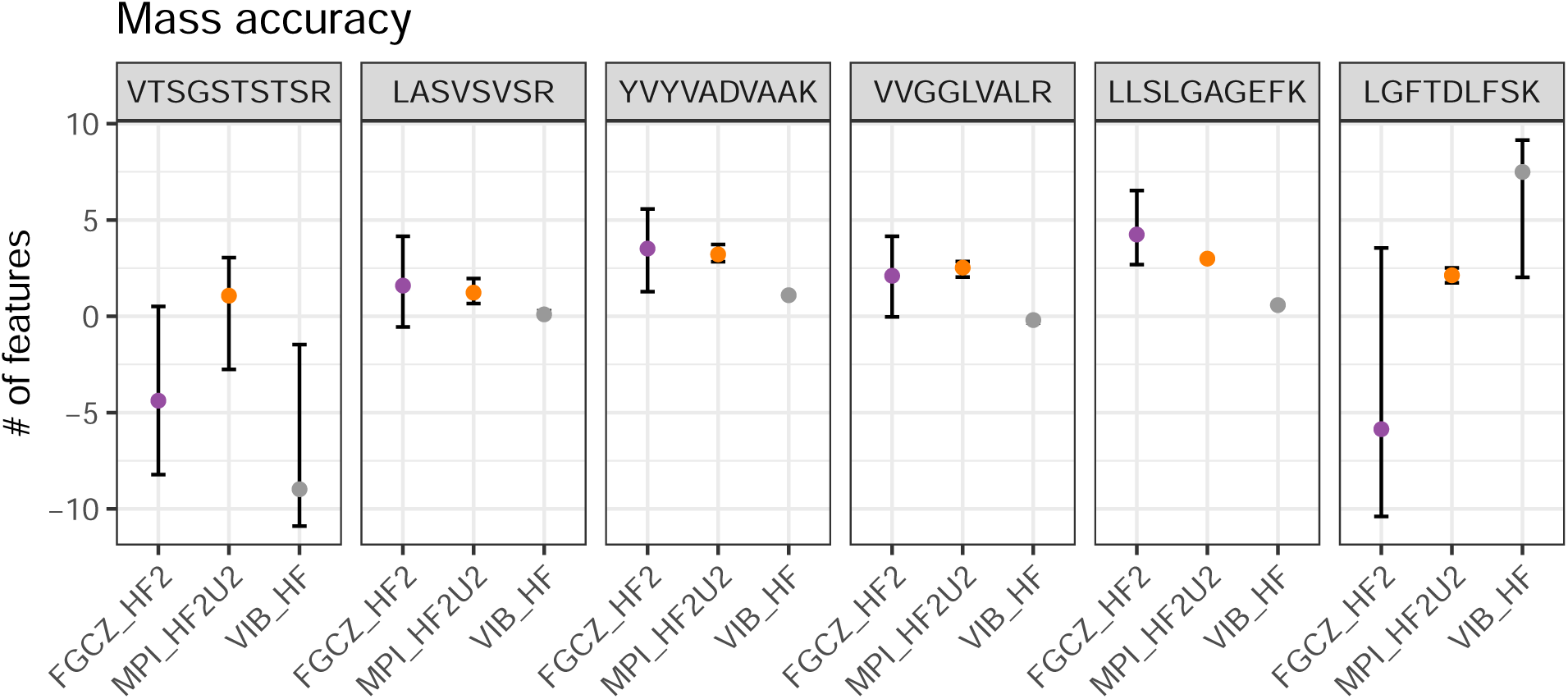

## Bibliography

Beri J, Rosenblatt MM, Strauss E, Urh M & Bereman MS (2015) Reagent for Evaluating Liquid Chromatography-Tandem Mass Spectrometry (LC-MS/MS) Performance in Bottom-Up Proteomic Experiments. Anal Chem 87: 11635–11640

Chiva C, Mendes Maia T, Panse C, Stejskal K, Douché T, Matondo M, Loew D, Helm D, Rettel M, Mechtler K, et al (2021) Quality standards in proteomics research facilities: Common standards and quality procedures are essential for proteomics facilities and their users. EMBO Rep 22: e52626.

Chiva C, Olivella R, Borras E, Espadas G, Pastor O, Sole A & Sabido E (2018) QCloud: A cloud-based quality control system for mass spectrometry-based proteomics laboratories. PLoS ONE 13: e0189209–e0189209

Fortier I, Raina P, Van den Heuvel ER, Griffith LE, Craig C, Saliba M, Doiron D, Stolk RP, Knoppers BM, Ferretti V, et al (2017) Maelstrom Research guidelines for rigorous retrospective data harmonization. Int J Epidemiol 46: 103–105

Lippens S, D’Enfert C, Farkas L, Kehres A, Korn B, Morales M, Pepperkok R, Premvardhan L, Schlapbach R, Tiran A, et al (2019) One step ahead: Innovation in core facilities. EMBO Rep 20: e48017

Meder D, Morales M, Pepperkok R, Schlapbach R, Tiran A & Van Minnebruggen G (2016) Institutional core facilities: prerequisite for breakthroughs in the life sciences: Core facilities play an increasingly important role in biomedical research by providing scientists access to sophisticated technology and expertise. EMBO Rep 17: 1088–1093

Olivella R, Chiva C, Serret M, Mancera D, Cozzuto L, Hermoso A, Borràs E, Espadas G, Morales J, Pastor O, et al (2021) QCloud2: An Improved Cloud-based Quality-Control System for Mass-Spectrometry-based Proteomics Laboratories. J Proteome Res 20: 2010–2013.

Pichler P, Mazanek M, Dusberger F, Weilnböck L, Huber CG, Stingl C, Luider TM, Straube WL, Köcher T & Mechtler K (2012) SIMPATIQCO: a server-based software suite which facilitates monitoring the time course of LC-MS performance metrics on Orbitrap instruments. J Proteome Res 11: 5540–5547

Trachsel C, Panse C, Kockmann T, Wolski WE, Grossmann J & Schlapbach R (2018) rawDiag: An R Package Supporting Rational LC-MS Method Optimization for Bottom-up Proteomics. J Proteome Res 17: 2908–2914.

Vizcaíno, J. A.; Deutsch, E. W.; Wang, R.; Csordas, A.; Reisinger, F.; Ríos, D.; Dianes, J. A.; Sun, Z.; Farrah, T.; Bandeira, N.; Binz, P.-A.; Xenarios, I.; Eisenacher, M.; Mayer, G.; Gatto, L.; Campos, A.; Chalkley, R. J.; Kraus, H.-J.; Albar, J. P.; Martinez-Bartolomé, S.; Apweiler, R.; Omenn, G. S.; Martens, L.; Jones, A. R.; Hermjakob, H. ProteomeXchange Provides Globally Coordinated Proteomics Data Submission and Dissemination. Nat. Biotechnol. 2014, 32 (3), 223–226.

